# Molecular basis of tRNA substrate recognition and modification by the atypical SPOUT methyltransferase Trm10

**DOI:** 10.64898/2025.12.12.694034

**Authors:** Suparno Nandi, Sarah E. Strassler, Debayan Dey, Catherine B. Whitney, Olivia B. Roumaya, Aiswarya Krishnamohan, George M. Harris, Lindsay R. Comstock, Jane E. Jackman, Graeme L. Conn

## Abstract

The evolutionarily conserved methyltransferase Trm10 catalyzes N1 methylation of guanosine 9 (G9) in select tRNAs, but the basis of specific substrate recognition and modification has remained unclear. Using an S-adenosyl-L-methionine analog, we trapped a post-catalytic state of the Trm10-tRNA^Gly^ complex for structure determination by cryogenic electron microscopy. Three distinct complexes were captured: two monomeric Trm10-tRNA complexes with distinct tRNA acceptor stem orientations (“closed” and “open”), and a minor dimeric complex with two Trm10s bound to the same tRNA. The monomeric structures identify conserved residues involved in tRNA interactions across a positively charged surface that guide G9 into the catalytic site and stabilize the flipped nucleotide. In the tRNA_open_ conformation, acceptor stem rotation weakens tRNA-protein contacts, consistent with a product-release state. The dimeric complex, supported by tRNA-dependent protein crosslinking and molecular dynamics (MD) simulations, positions one Trm10 on G9 similarly to the monomeric complexes, while the other contacts distal tRNA regions, suggesting a functional role in promoting functionally critical conformational transition(s). MD simulations also show how Trm10 achieves selective stabilization of G9 over A9 in the binding pocket. Overall, our findings reveal the mechanism of G9-specific tRNA methylation by Trm10 and suggest a unique mechanism of action among RNA-modifying SPOUT methyltransferases.

## INTRODUCTION

Methylation is one of the most common chemical modifications in biology and plays important roles in gene expression, small molecule metabolism, and regulation of macromolecule structure and function (1–4). The SpoU-TrmD (SPOUT) enzyme family–one of five major classifications of S-adenosyl-L-methionine (SAM)-dependent methyltransferases–was designated upon identification of structural similarity between the transfer RNA (tRNA)-modifying enzymes TrmH (also known as SpoU) and TrmD (5–10). SPOUT (or Class IV) methyltransferases are characterized by a unique α/β fold and a deep trefoil knot in the C-terminal half of the catalytic domain. This protein knot is functionally important as it stabilizes SAM in a bent conformation necessary for methyl transfer by this family of enzymes (7,11–13).

The SPOUT tRNA methyltransferase Trm10 modifies the N1 base position of the 9^th^ nucleotide in the core region of some tRNAs and is evolutionarily conserved across Eukarya and in many Archaea (14). *Saccharomyces cerevisiae* Trm10 was first predicted to be a member of the SPOUT family in 2007 (9), and this designation was experimentally verified through X-ray crystallographic studies of the C-terminal SPOUT domains of Trm10 from *S. cerevisiae* and *Schizosaccharomyces pombe* (15). In *S. cerevisiae*, Trm10 methylates G9 of 13 of 26 possible tRNA substrates (16). In contrast, Trm10 enzymes in the Archaea *Sulfolobus acidocaldarius* and *Thermococcus kodakarensis* were found to either methylate A9 or exhibit dual G9/A9 activity, respectively (17–19). Humans express three Trm10 enzymes which also possess diversity in their substrate specificities and localization. TRMT10A (the direct homolog of yeast Trm10), which methylates many but not all tRNAs with G9, and TRMT10B, which methylates A9 in tRNA^Asp^, are both cytosolic enzymes, while the mitochondria-localized TRMT10C modifies both G9 and A9 of all tRNAs that contain a purine at this position (16,20–22). Among the human enzymes, the biological importance of TRMT10A has been highlighted by studies linking loss of its activity to neurological and endocrine disorders (23–27). Similarly, in *S. cerevisiae*, loss of Trm10 leads to a variety of detrimental effects, such as increased susceptibility to 5-fluorouracil (28), compromised tRNA integrity (29), and decreased translational fidelity (30).

The structures of the yeast Trm10 catalytic C-terminal domain (CTD) confirmed the presence of a typical knotted SPOUT methyltransferase fold (15). However, unlike all other known RNA-modifying SPOUT enzymes which function as homodimers, Trm10 was observed as a monomer in the crystal structure. Subsequent structures of archaeal Trm10 enzymes from *S. acidocaldarius* and *T. kodakarensis* revealed that these enzymes also lack the expected dimeric SPOUT methyltransferase structure (17,31). Identification that Trm10 functions as a monomeric enzyme was unexpected because the catalytic centers of other SPOUT RNA methyltransferases are formed at the dimer interface, with both protomers making essential contributions to the other’s active site. Currently, only one other monomeric SPOUT methyltransferase, Sfm1, has been identified but this enzyme is also unique in other ways, such as acting on a protein substrate and containing a distinct negatively-charged surface surrounding the active site (32), compared to the positive surface of RNA-modifying SPOUT enzymes. As such, it remains unclear how Trm10 recognizes and efficiently methylates its tRNA substrate without dimerization.

More recently, the first structures of a Trm10 enzyme (human TRMT10C) bound to substrate tRNA were determined as part of the mitochondrial protein-only RNase P (PRORP) complex (33,34), and have begun to address this question albeit within the context of multiple additional protein binding partners. Within the PRORP complex, TRMT10C serves as one of the enzyme subunits responsible for pre-tRNA recognition, modification, and processing. These structures show the NTD of TRMT10C wrapping around the tRNA so that the substrate is enclosed on each side by the catalytic SPOUT CTD and the NTD, with the target A9 base flipped into the enzyme active site. However, other subunits of the RNase P complex also contact the pre-tRNA, including the pentatricopeptide repeat and nuclease, which have known roles in RNA recognition and are necessary for tRNA 5’-end cleavage by the nuclease subunit. Importantly, TRMT10C is the only SPOUT methyltransferase that requires participation in a larger complex for its enzymatic activity which may explain why it is able to function as a “monomer”. In contrast, other Trm10 enzymes do not require accessory factors for methyltransferase activity, and it therefore remains unclear how yeast Trm10 and its direct human homolog TRMT10A are able to recognize their correct tRNA substrate(s) as atypical stand-alone monomeric SPOUT methyltransferases.

Here, we use single-particle cryogenic electron microscopy (cryo-EM) to gain structural insight into the mechanism of tRNA^Gly-GCC^ substrate recognition by *S. cerevisiae* Trm10. Multiple states of the Trm10-tRNA complex were identified, including two “monomeric” Trm10-tRNA complexes, distinguished primarily by the orientation of the tRNA acceptor stem, and a minor population with two Trm10 proteins on a single tRNA ((Trm10)_2_-tRNA), which was corroborated by observation of tRNA-dependent recruitment of two Trm10 proteins in crosslinking experiments. The three different Trm10-tRNA complex structures, supported by molecular dynamics (MD) simulations, reveal key interactions that are critical for substrate tRNA recognition and G9 modification, elucidating the molecular determinants of guanosine selectivity, and suggesting a new mechanistic model for Trm10 function as an atypical monomeric SPOUT methyltransferase.

## MATERIAL AND METHODS

### Trm10 expression and purification

Full-length wild-type Trm10 from *S. cerevisiae* with an N-terminal 6xHis-tag was expressed from the pET-derived plasmid pJEJ12-3 in *E. coli* BL21(DE3)-pLysS grown in lysogeny broth, as described previously (14). Briefly, protein expression was induced by the addition of 1 mM β-D-1-thiogalactopyranoside at mid-log phase growth (OD_600_ ∼0.6), and growth continued at 37°C for an additional 5 hours. All steps during lysis and initial purification were performed in 20 mM HEPES pH 7.5, 4 mM MgCl_2_, 1.0 mM β-mercaptoethanol (BME), 10 mM imidazole, and 5% (v/v) glycerol. To ensure removal of co-purifying SAM, cells were lysed in this buffer with 1 M NaCl and 0.5% (v/v) Triton X-100 added, and the lysate was dialyzed three times against the same buffer but containing 2 M NaCl. A final dialysis step was used to reduce the NaCl to 0.25 M for protein purification by sequential Ni^2+^-affinity (HisTrap HP), heparin-affinity (HiPrep Heparin 16/10, GE Healthcare), and gel filtration (Superdex 75 16/600, GE Healthcare) chromatographies on an ÄKTApurifier10 system (GE Healthcare). Trm10 was eluted from the gel filtration column in 20 mM Tris (pH 7.5) buffer containing 100 mM NaCl, 1 mM MgCl_2_, 5 mM BME, and 5% (v/v) glycerol and flash frozen in liquid nitrogen before storage at −80°C.

Trm10 N-terminal domain (NTD) truncation and multi-site variants were constructed using Phusion mutagenesis (Thermo Fisher Scientific), and sequence-validated constructs were used for expression in *E. coli* BL21(DE3), as described for wild-type Trm10. NTD variant enzymes (Trm10-Δ47, Trm10-Δ63 and Trm10-8A) used for *in vitro* methyltransferase activity and tRNA binding assays were purified using a single Talon resin (Takara Bio) purification step as described previously (16).

### RNA *in vitro* transcription and purification

tRNA^Gly-GCC^ was *in vitro* transcribed from *BstNI* linearized plasmid DNA using T7 RNA polymerase as previously described (35). Briefly, *in vitro* transcription was performed for 5 hours at 37°C in 200 mM HEPES-KOH (pH 7.5) buffer containing 28 mM MgCl_2_, 2 mM spermidine, 40 mM dithiothreitol (DTT), 6 mM each rNTP, and 100 µg/mL DNA template. Following addition of 40 mM EDTA to clear pyrophosphate-magnesium precipitates and dialysis against TE buffer (10 mM Tris-HCl, 1 mM EDTA, pH 8.0), tRNA^Gly-GCC^ was purified by denaturing polyacrylamide gel electrophoresis (8.3 M urea, Tris-Borate-EDTA buffer) and eluted from the gel using the crush and soak method in 0.3 M sodium acetate, followed by ethanol precipitation, as previously described (35).

### Trm10-tRNA complex formation and cryo-EM specimen preparation

The SAM analog 5’-(diaminobutyric acid)-N-iodoethyl-5’-deoxyadenosine ammonium hydrochloride (“NM6”) was prepared as previously described (36), building upon earlier methyltransferase-directed aziridinium cofactor chemistry (37), and purified by semi-preparative reverse-phase HPLC. *In situ* activation of NM6 results in a Trm10 cosubstrate that is covalently attached by the enzyme to tRNA (**Supplementary Fig. S1**) (38,39). Prior to the preparation of the Trm10-tRNA complex for cryo-EM, glycerol was removed from the Trm10 sample by dialysis against 20 mM Tris buffer (pH 7.5) containing 100 mM NaCl, 1 mM MgCl_2_, and 5 mM BME. NM6 was dissolved in the same buffer. tRNA^Gly-GCC^ was incubated at 65°C in TE buffer for 10 minutes and then slowly cooled to room temperature. The Trm10-tRNA-NM6 complex was formed by mixing Trm10 (60 µM), tRNA (30 µM), and NM6 (600 µM) in a 2:1:20 molar ratio, followed by incubation at 30°C for 30 minutes. The sample was diluted in the same buffer as used for Trm10 and NM6 to a complex concentration of 2.25 µM or 4.5 µM. The diluted complex (3 µl) was applied to freshly glow-discharged grids (UltrAufoil R 0.6/1, 300 Mesh, Electron Microscopy Sciences), with blotting for 2 or 3 s at 100% humidity at room temperature before freezing in liquid ethane using a Vitrobot Mark IV System (Thermo Scientific). Grids were stored in liquid nitrogen until used for data collection.

### Cryo-EM image collection, processing and analysis

Data were collected on a Titan Krios microscope (FEI) operating at 300 keV with a K3 direct electron detector (Gatan) at the National Center for Cryo-EM Access and Training (NCCAT). A total of 26,656 micrographs were collected with a defocus range of −0.8 to −2.0 μm at 105,000x magnification with a 0.412 Å/pixel size. The dataset contains micrographs that were collected from tilting the sample 0°, 30°, and 45° to capture the complex in a wider distribution of orientations. Micrographs were collected as 50 frames with a dose rate of 29.91 e^-^/Å^2^/s and a total exposure of 2.0 seconds, for an accumulated dose of 59.82 e^-^/Å^2^.

The complete workflow for cryo-EM structure determination of the three structures in CryoSPARC (40) is summarized in **Supplementary Fig. S2**. First, the micrographs were imported and motion corrected by Patch Motion Correction, followed by estimation of contrast transfer function (CTF) parameters using Patch CTF. Blob Picker was used for particle picking, and incorrectly selected particles were discarded after reference-free two-dimensional (2D) class averaging. The best 2D classes were used for template-based particle picking, followed by multiple rounds of reference-free 2D class averaging, resulting in 5,634,445 and 244,276 particles (256-pixel box size) corresponding to monomeric Trm10-tRNA and dimeric (Trm10)_2_-tRNA complexes, respectively. Multiple *ab-initio* 3D reconstructions were generated from these particles with C1 symmetry, and good-quality maps with observable features for both Trm10 and tRNA were used as a reference for 3D heterogeneous refinement. For the (Trm10)_2_-tRNA complex, heterogeneous refinement of *the ab-initio* map did not yield improved results, and the results were therefore discarded.

For both monomeric Trm10-tRNA complexes, the best maps from heterogeneous refinement were used for non-uniform (NU) refinement followed by CTF refinement, reference-based motion correction, and further NU refinement. This process yielded a final global map resolution of 3.63 Å based on gold-standard refinement Fourier Shell Correlation (0.143 cutoff) for the tRNA_closed_ conformation (**Supplementary Fig. S3A**). For the tRNA_open_ conformation, the resultant NU refined map was 3D classified, and the best classes were chosen for further NU refinement, resulting in a final global map resolution of 3.37 Å (**Supplementary Fig. S3B**).

For the dimeric (Trm10)_2_-tRNA structure, NU refinement was performed with the best maps from the *ab-initio* refinement, followed by reference-based motion correction and 3D classification. The best classes were chosen for NU refinement, generating a final map of 3.89 Å resolution (**Supplementary Fig. S3C**). Finally, the NU refined maps from the three structures were sharpened in CryoSPARC, and the half-maps were sharpened using DeepEMhancer (41). CryoSPARC local resolution and orientation diagnostics tools were used to generate local resolution maps and Sampling Compensation Factor values, respectively, for all maps (**Supplementary Fig. S3D-I**).

### Model building and refinement

The NU refined maps were used for initial model building of the three Trm10-tRNA complexes. A yeast Trm10 structure (PDB code 4JWJ) (15) with ligands and water molecules removed was docked in the maps for the open and closed conformations, while two copies of the yeast Trm10 structure were used for the (Trm10)_2_-tRNA map. As there is no available structure of tRNA^Gly-GCC^, we generated a 3D model using AlphaFold 3 (42) with the sequence of tRNA^Gly-GCC^. A single molecule of the tRNA^Gly-GCC^ was docked into each of the three maps using Chimera (v1.17) (43). The combined model was refined in PHENIX (44) using rigid body, global minimization, simulated annealing, local grid search, and B-factor refinement. Manual adjustment of the refined model and de-novo modeling of NM6 modified-G9 was done in COOT (45) using the DeepEMhancer (41)- and CryoSPARC-sharpened maps, followed by a B-factor refinement in PHENIX. All three structures were validated using PHENIX. Comprehensive details on data collection, processing, model construction, refinement, and validation are provided in **Table 1**.

**Table 1:** Cryo-EM data collection, refinement, and model validation for the Trm10-tRNA complexes.

|  | Trm10-tRNA<sub>closed</sub> | Trm10-tRNA<sub>open</sub> | (Trm10)<sub>2</sub>-tRNA |
| --- | --- | --- | --- |
| **Deposition** |  |  |  |
| Coordinates (PDB) | 9XZQ | 9XZR | 9XZS |
| Map (EMDB) | EMD-72368 | EMD-72369 | EMD-72370 |
| **Data collection and processing** |  |  |  |
| Microscope | TFS Titan Krios | TFS Titan Krios | TFS Titan Krios |
| Camera | Gatan K3 | Gatan K3 | Gatan K3 |
| Voltage, kV | 300 | 300 | 300 |
| Magnification (×) | 105,500 | 105,500 | 105,500 |
| Electron exposure, e<sup>-</sup>/Å<sup>2</sup> | 59.82 | 59.82 | 59.82 |
| Defocus range, μm | -0.8 to -2.0 | -0.8 to -2.0 | -0.8 to -2.0 |
| Pixel size, Å | 0.412 | 0.412 | 0.412 |
| Symmetry | C1 | C1 | C1 |
| No. particles, initial | 5,634,445 | 5,634,445 | 244,276 |
| final | 1,398,183 | 1,440,392 | 156,897 |
| Map resolution (FSC 0.143), Å | 3.63 | 3.37 | 3.89 |
| **Refinement and model** |  |  |  |
| Model resolution (FSC 0.143, unmasked), Å | 3.9 | 3.4 | 4.4 |
| CC<sub>mask</sub> | 0.76 | 0.73 | 0.54 |
| Non-hydrogen atoms | 3090 | 2981 | 4513 |
| Protein residues | 189 | 177 | 360 |
| RNA residues | 71 | 71 | 71 |
| SAM analog NM6 (AN6 “ligand”) | 1 | 1 | 1 |
| B factors (min./ max./ mean), Å<sup>2</sup> |  |  |  |
| Protein | 170.4/399.5/252.9 | 94.4/299.3/176.8 | 100.8/364.0/225.0 |
| RNA | 202.1/458.0/262.8 | 154.4/357.2/213.5 | 182.4/1016.1/359.8 |
| SAM analog NM6 (AN6 “ligand”) | 249.8/249.8/249.8 | 154.4/154.4/154.4 | 190.1/190.1/190.1 |
| RMS deviations |  |  |  |
| Bond lengths, Å | 0.016 | 0.013 | 0.011 |
| Bond angles, ° | 1.643 | 1.402 | 1.271 |
| **Validation** |  |  |  |
| MolProbity score | 1.87 | 2.57 | 2.40 |
| Clashscore | 8.67 | 12.9 | 9.7 |
| Rotamer outliers, % | 2.3 | 3.1 | 2.4 |
| Ramachandran plot (protein) |  |  |  |
| Favored, % | 97.3 | 89.1 | 87.9 |
| Allowed, % | 2.7 | 10.9 | 11.3 |
| Disallowed, % | 0.0 | 0.0 | 0.8 |

Structural figures were prepared using PyMOL (v3.1) and Chimera X (v1.9) (46,47), using the DeepEMhancer- and the CryoSPARC-sharpened maps. Modevectors representing the normal mode analysis of the structural transition in the tRNA conformations were obtained using the *modevectors.py* script available from PyMOLWiki (48). Buried surface area calculations were performed using the PDBePISA webserver (v1.52) (49). Average residue interaction energy, which includes nonbonded electrostatic and van der Waals terms, was calculated over all residue pairs (protein–protein, RNA–RNA, and protein–RNA) in Maestro (BioLuminate module, Schrödinger 2024-3) using the OPLS4 force field.

### Nuclease (RNase T1) footprinting

^32^P-5’-end labeled *in vitro* transcribed tRNA was subjected to partial degradation with RNase T1 (0.01 U/ml; Ambion) either in the absence or presence of Trm10 (at 1 or 5 µM final concentration) for 1 hour at 37 °C. Reactions were stopped by the addition of phenol:chloroform:isoamyl alcohol (25:24:1) and tRNA fragments were purified by phenol extraction and ethanol precipitation, and run on a 10% polyacrylamide denaturing (8M urea) gel. Gels were dried and exposed to a phosphor screen and scanned using TyphoonTM imaging system (GE Healthcare) and quantified using ImageQuantTM TL software (GE Healthcare). All reactions were done in duplicate and run on the same gel to reduce variability. Alkaline hydrolysis ladders were prepared by incubating the labeled tRNA in 50 mM sodium carbonate for 10 minutes at 75 °C and stopped by the addition of denaturing gel loading buffer.

### Trm10*-*tRNA binding

Assays to measure tRNA binding were performed with 5’-6-carboxyfluorescein-labeled tRNA^Gly-GCC^ using fluorescence anisotropy as previously described (38). Anisotropy values (FA) at each concentration (5-300 nM) of Trm10 (WT or variant, [E]) were measured in triplicate independent experiments, each comprising three technical replicates. The resulting independent replicate data were individually fit to the binding isotherm (Eq. 1) to determine the observed K_D,app_ and n_H_ (Hill coefficient) for each variant and associated error of the fit:

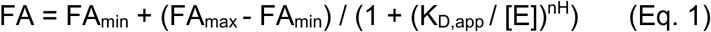

The resulting three independent measurements for each protein were then averaged to yield the reported K_D,app_. Reported errors were generated by propagation of error from the n=3 experimental values determined for K_D,app_.

### Trm10 *in vitro* methyltransferase activity

The Trm10 substrate tRNA^Gly-GCC^ was *in vitro* transcribed as described above, except that the tRNA product was uniformly radiolabeled at all 22 guanosine nucleotides with ^32^P by the inclusion of [α^32^P]-GTP, as described previously (16). After denaturing gel purification of the labeled tRNA, *in vitro* methylation assays were performed using the indicated enzymes, labeled tRNA (2000 cpm per reaction) and 0.5 mM SAM. Following a two-hour incubation at 30 °C, reactions were quenched by addition of phenol:chloroform:isoamyl alcohol (25:24:1, v:v:v), and tRNA recovered by ethanol precipitation and digested with nuclease P1 (Sigma-Aldrich), under conditions described previously (20), to generate 5’-^32^P-nucleotides (either p*m^1^G9 after Trm10 modification, or p*G from unmethylated G nucleotides throughout the tRNA). Free labeled nucleotides were resolved by cellulose TLC in isobutyric acid:H_2_O:NH_4_OH (66:33:1, v:v:v) solvent and imaged with a Typhoon imager (Cytiva). Percent modification at position 9 was quantified by calculating the percent of p*m^1^G/p*G signal at each enzyme concentration and multiplying by 22 to correct for the 22 total G nucleotides that were labeled throughout the tRNA, such that 100% modification of G9 corresponds to ∼4.5% of total labeled G residues (1/22) converted to p*m^1^G9 product.

### Bis(sulfosuccinimidyl)suberate (BS3) protein crosslinking

Wild-type or variant Trm10 and tRNA^Gly-GCC^ were dialyzed against 20 mM HEPES pH 7.5 buffer containing 150 mM NaCl, and 5 mM BME. Immediately before use, BS3 crosslinking reagent (Thermo Scientific Pierce BS^3^, #21580) was prepared in water to a final concentration of 12.5 mM and added in a 50-fold molar excess to samples containing 6 μM Trm10 in the presence or absence of 2 μM tRNA^Gly-GCC^ and SAM. Samples were incubated at room temperature for 30 minutes and quenched with Tris pH 7.5 to a final concentration of 50 mM before analysis on a 9% polyacrylamide SDS-PAGE gel and visualization by Coomassie blue staining.

### Structural modeling of target nucleotide interactions and MD simulations of Trm10-tRNA complexes

To investigate nucleotide specificity at position 9 (G9 vs. A9) in Trm10, a monomeric, Trm10-tRNA_open_ complex with a G9 to A9 mutation was made using the Maestro mutagenesis tool of the Schrödinger software suite (version 2024-4), and the resulting variant structure (Trm10-tRNA^A9^) was energy minimized to relieve steric clashes and optimize geometry. Wild-type and tRNA mutant complexes included tRNA nucleotides 1 to 71 and Trm10 residues 87 to 262 (i.e., excluding the disordered NTD), and NM6 was replaced with SAM in its established binding site from a previously reported structure (PDB 4JWJ) (15).

Structural models of dimeric Trm10–tRNA complexes were also similarly generated in the Schrödinger suite (version 2024-4) to probe the contribution of protein–tRNA interface residues to the binding of each Trm10 protomer. For each complex, Trm10 variants with six alanine substitutions (“6A”: R121A, R127A, R128A, N154A, N156A, and N159A) were generated. The 6A substitutions were made in both Trm10 protomers, as well as individually in each of the catalytic (Trm10) and auxiliary (Trm10’) protomers for a total of three distinct systems for simulation. All complexes included tRNA nucleotides 1–71 and Trm10 residues 94–264 for the Trm10 protomer (excluding the disordered N-terminal domain) and 87–274 for Trm10’. NM6 was replaced with SAM, and all systems were subjected to energy minimization, as before.

Each system was prepared for MD using the Protein Preparation Wizard in Maestro to assign bond orders, add hydrogens, and set protonation states appropriate for pH 7.5 using Epik. MD simulations were performed using the OPLS4 force field and solvation in an orthorhombic TIP3P water box, with system neutralization at physiological ionic strength accomplished with 150 mM NaCl. System relaxation followed the default multistep relaxation protocol in Desmond, including restrained minimization, restrained and unrestrained equilibration stages, and gradual heating from 0 K to 310.15 K. The final equilibration phase was carried out for 10 ns under NPT conditions at 310.15 K and 1 atm pressure, using the Nose-Hoover thermostat and Martyna– Tobias-Klein barostat. For each system, production simulations were run for 100 ns in the NPT ensemble, with temperature maintained at 310.15 K and pressure at 1 atm. Three independent replicates were performed with unique initial velocities, and coordinates were saved every 100 ps for analysis.

Trajectory analyses were conducted using Desmond tools in the Schrödinger suite, including calculation of root mean square deviation (RMSD) using to assess overall complex structural variability and root mean square fluctuation (RMSF) to quantify residue-level flexibility. RSMD and RSMF were both calculated based on backbone atom positions (C_α_ or P). In addition, energy component analyses were performed, including total and decomposed interaction energies, with comparisons carried out at both chain and residue levels to evaluate the effects of 6A substitutions and conformational states on Trm10–tRNA interaction.

### Trm10 protein family sequence analysis

The Trm10 protein sequence from *S. cerevisiae* was used as a query to search for tRNA methyltransferase Trm10-type domain protein family members in fungi in the UniProtKB database. The search yielded 1687 fungal sequences for which Uniref50 IDs were assigned. After removing redundant entries based on UniRef50 clustering, 321 unique fungal sequences were retained for multiple sequence alignment in Geneious Prime using the Blosum62 matrix, followed by calculation of residue identity.

## RESULTS

### Structure of the *S. cerevisiae* Trm10-tRNA^Gly-GCC^ complex

To determine the cryo-EM structure of the Trm10-tRNA^Gly-GCC^ complex, we used a SAM analog (“NM6”) that is transferred and covalently attached to the G9 target nucleotide in its entirety, trapping Trm10 on its tRNA substrate (**Supplementary Figs. S1-S3**) (35). Our use of NM6 was motivated by prior observations that Trm10 can bind with similar affinity to both substrate and non-substrate tRNAs (38), and our finding that only in the presence of active Trm10 and NM6 was a unique protein-tRNA complex observed via native gel electrophoresis (**Supplementary Fig. S1B**). Most importantly, the complex trapped via NM6 has undergone G9 modification by Trm10, ensuring that all particles containing Trm10, NM6, and tRNA represent a *bone fide* state during the process of tRNA modification. As Trm10 is trapped after transfer of NM6 to the tRNA G9 nucleotide, we refer to this complex as an immediately post-catalytic state.

Trm10 is bound to tRNA^Gly-GCC^ in a predominantly 1:1 ratio in two distinct monomeric complexes that are distinguished by the orientation of the tRNA acceptor stem (“open” or “closed”, tRNA_open_ and tRNA_closed_, respectively; **Fig. 1A,B**). In both structures, only the Trm10 SPOUT domain (starting with residue N87; **Supplementary Fig. S4**), tRNA, and NM6 are well defined, with no map corresponding to the Trm10 NTD, which is therefore absent from both models. The structure of a third complex containing two Trm10 SPOUT domains, (Trm10)_2_-tRNA^Gly-GCC^, was also determined from a minor fraction of the particles (discussed further below). Again, no well-defined map was observed for either Trm10 NTD.

**Fig. 1.**
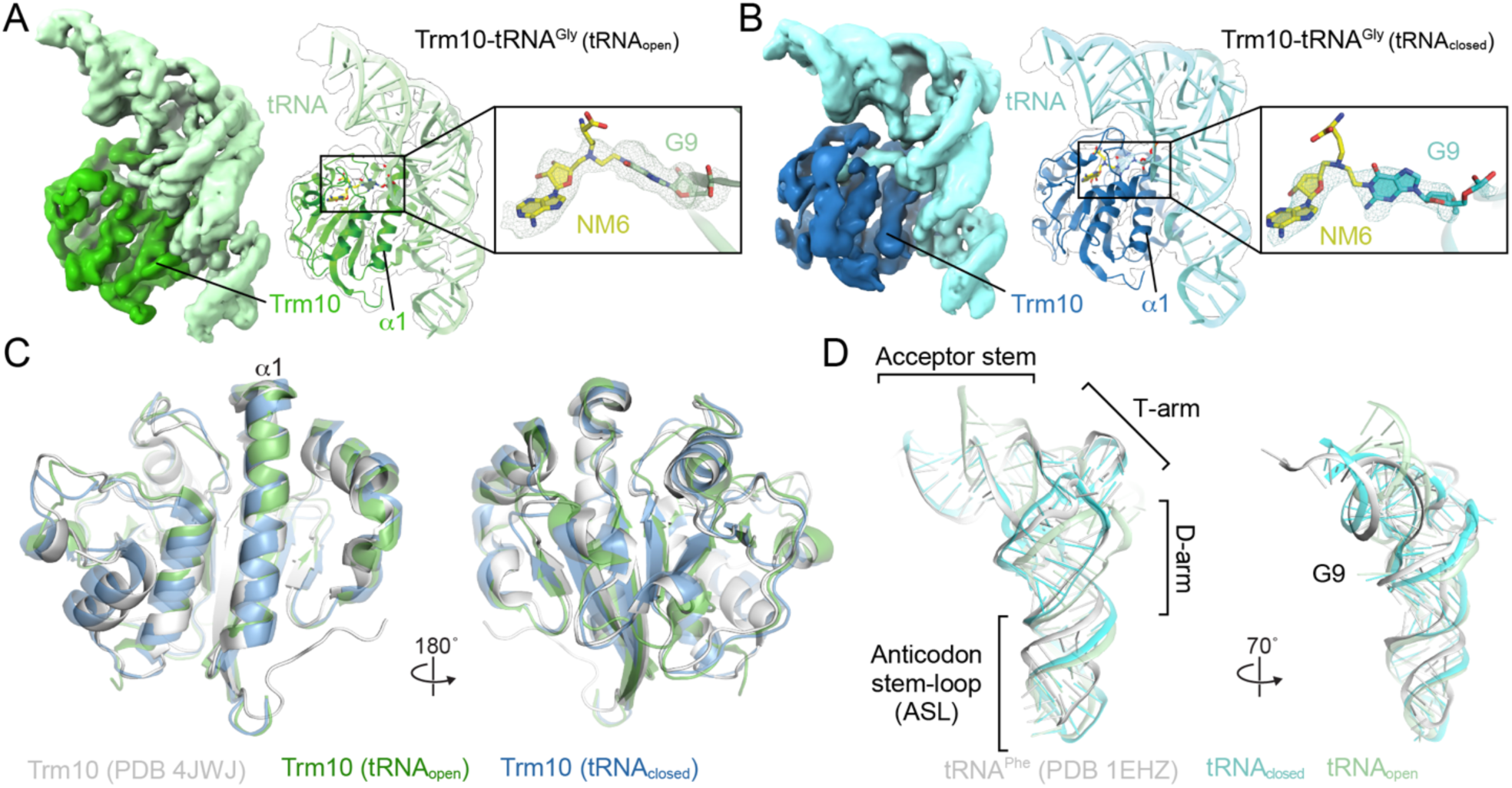
Structure of the monomeric Trm10-tRNA^Gly^ complex. ***A,*** DeepEMhancer (DEM)-sharpened map of the Trm10-tRNA^Gly^ complex with the tRNA_open_ conformation at 3.37 Å resolution (Trm10 and tRNA are shown in green and light green, respectively). The final model is shown within a semi-transparent white map outline. ***Inset***, Zoomed-in view of NM6 (yellow) covalently linked to G9 within DEM-sharpened map shown (threshold: 0.234; value range: −0.00338 to 2.19). ***B,*** As for panel A, but for the monomeric Trm10-tRNA complex with the tRNA_closed_ conformation at 3.63 Å resolution (Trm10 and tRNA shown in blue and cyan, respectively). For the *inset* in *panel B*, the map is DEM-sharpened with a threshold of 0.591 and a map value range of −0.00171 to 2. ***C,*** Alignment of Trm10 from both monomeric tRNA complexes and free Trm10 (white) shows that the overall structure remains largely unchanged. ***D,*** Superimposition of tRNA_closed_, tRNA_open_, and free tRNA^Phe^ (PDB 1EHZ) (50) reveals significant differences throughout the molecule, particularly in tRNA_open_.

In both monomeric complexes, Trm10 is structurally similar to the previously reported yeast Trm10 SPOUT domain structure (PDB code 4JWJ; **Fig. 1C**)(15), with root mean square deviation (RMSD) for alignment of 1.5 Å (with tRNA_open_) and 1.4 Å (with tRNA_closed_) over 159 and 186 C_α_ atoms, respectively. The positioning of Trm10 on the tRNA is also consistent with the position of the Trm10 paralog TRMT10C in relation to its tRNA substrate as part of the mitochondrial RNase P complex, further supporting the relevance of these structures for substrate recognition by Trm10 (**Supplementary Fig. S5A**) (33). Trm10 makes extensive contacts with the core of the tRNA and the D-arm, with its α1 helix interacting directly with G9, which is covalently linked to NM6 (**Fig. 1A-C**). In contrast, Trm10 makes more limited contact with the tRNA anticodon stem-loop (ASL), which has a poorer map quality compared to the rest of the structure (**Supplementary Fig. S3D,E**) and appears to be more dynamic during substrate recognition by Trm10, with the highest B-factors observed for the loop nucleotides (**Supplementary Fig. S5B**).

The absence of map corresponding to the Trm10 NTD (**Supplementary Fig. S4**) was unexpected as this domain was previously proposed to be functionally important, with complete deletion of the NTD (residues 1-83) resulting in a substantial reduction in activity (15). We tested three additional Trm10 variants with shorter NTD truncations of 47 (Trm10-Δ47) or 63 amino acids (Trm10-Δ63), and with substitution of 8 clustered Arg/Lys residues to Ala within residues 47-64 of the NTD (Trm10-8A); **Supplementary Fig. S6A**). In contrast to the larger truncation, each new variant retained essentially wild-type binding affinity for tRNA^Gly-GCC^, with only a ∼2.5-fold reduction observed for the most impacted variant, Trm10-8A (**Supplementary Fig. S6B**). Further, despite its modestly reduced tRNA binding affinity, Trm10-8A retained near wild-type level activity, while the truncated Trm10-Δ47 and Trm10-Δ64 variants showed intermediate and essentially no activity, respectively (**Supplementary Fig. S6C**). For example, at equivalent concentrations (0.01 µM), m^1^G9 formation for Trm10-8A, Trm10-Δ47, and Trm10-Δ63 were approximately 85%, 13%, and 2% of wild-type (100%), respectively (lanes indicated with arrowheads in **Supplementary Fig. S6C**). Collectively, these results indicate that the majority of the Trm10 NTD is dispensable for tRNA binding, and the Trm10-tRNA contacts observed in our structures are therefore likely the major drivers of complex formation. However, a shorter region of the NTD, proximal to the SPOUT domain and likely located near the tRNA anti-codon, appears critical for the modification process itself.

We speculate that the short functionally important NTD region may engage with the tRNA ASL in an essential step prior to methylation and is thus not observed in the post-catalytic structure captured using the NM6 analog. Dissociation of this region after modification is also consistent with the higher flexibility observed for the tRNA ASL. To explore potential Trm10 NTD-tRNA interaction prior to modification, we generated and aligned an AlphaFold (42) model of full-length Trm10 to the Trm10 SPOUT domain in the tRNA_closed_ complex (RMSD of 1.4 Å, over 181 C_α_ atoms). This modeling shows the potential for the NTD to wrap around the tRNA in a binding mode similar to the TRMT10C-tRNA complex (33), positioning the NTD region proximal to the SPOUT domain adjacent to the tRNA ASL (**Supplementary Fig. S7A**). Additionally, calculation of the Trm10 electrostatic surface potential reveals that this modeled NTD position places a second positively charged surface against the opposite side of the tRNA from that of the CTD (**Supplementary Fig. S7B**), which could favor complex formation but does not appear essential given the retained activity of the Trm10-8A and Trm10-Δ47 variants.

### Trm10 induces structural changes throughout the bound tRNA

Superimposition of tRNA^Gly-GCC^ from the tRNA_open_ and tRNA_closed_ monomeric Trm10 complexes with tRNA^Phe^ (which serves as an established “standard” tRNA for which a high-resolution structure is available; PDB code 1EHZ) (50) yields RMSDs of 6.7 Å and 5.3 Å, respectively, over 68 phosphate atoms. Structural divergence between tRNA^Phe^ and tRNA_closed_/ tRNA_open_ arises from conformational changes in each of the four major tRNA domains: acceptor stem, D-arm, T-arm, and ASL (**Fig. 1D**). The ASLs in both tRNA_open_ and tRNA_closed_ are similar, suggesting that this region undergoes consistent structural adaptation with Trm10 bound.

To enable visualization of phosphate atom movements using modevector analysis (48), a “free” tRNA^Gly-GCC^ structure was generated using AlphaFold. Comparison of this free structure with tRNA_closed_ reveals shifts in phosphate positions primarily in the ASL as well as the D-arm (**Fig. 2A**, *left*). This observation is supported by nuclease footprinting of free and Trm10-bound tRNA^Gly-GCC^, which shows a strong Trm10-dependent increase in accessibility to RNase T1 at G34, consistent with significant distortion in the ASL (**Supplementary Fig. S8**). Modevector analysis comparing the free and tRNA_open_ structures reveals similar changes in the ASL to those in tRNA_closed_ as well as more extensive shifts in the D-arm and additional changes in the T-arm and acceptor stem (**Fig. 2A**, *center*). These differences suggest a large-scale conformational change is induced in tRNA_open_ by Trm10. Finally, comparison of tRNA_closed_ and tRNA_open_ confirms that the major differences in these structures arise from their distinct acceptor stem orientations and movements in the adjacent D- and T-arm regions (**Fig. 2A**, *right*).

**Fig. 2.**
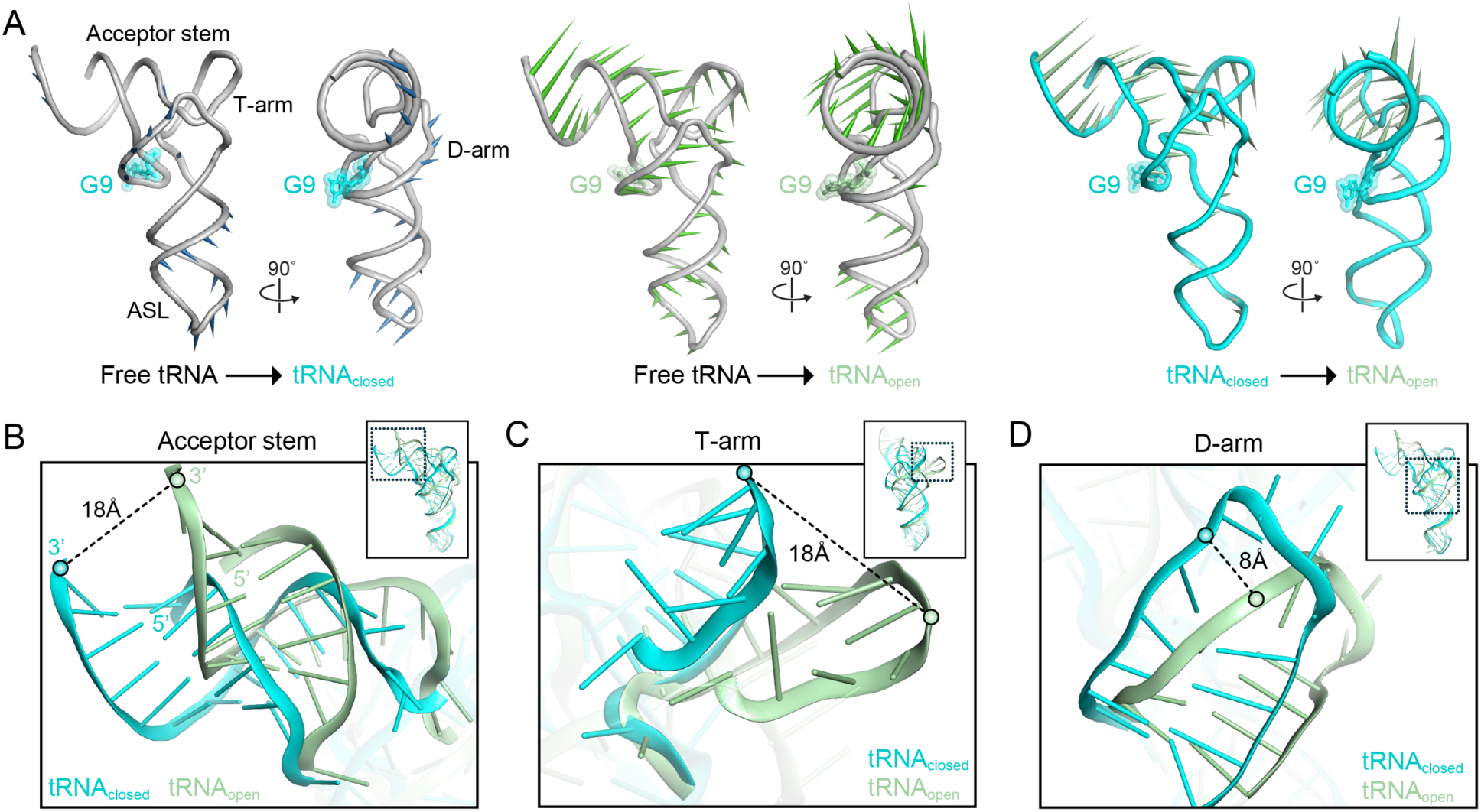
tRNA_open_ and tRNA_closed_ complexes with Trm10 exhibit distinct architectures. ***A,*** Modevector visualization of the tRNA conformational changes (P atom position) comparing (*left* to *right*): free tRNA *vs.* tRNA_closed_, free tRNA *vs.* tRNA_open_, and tRNA_closed_ *vs.* tRNA_open_. ***B,*** The acceptor stem of tRNA_open_ undergoes a rotation of 23° compared to tRNA_closed_, accompanied by an 18 Å displacement of its 3′ end. ***C,*** The T-arm rotates by 41° and undergoes a maximum shift of 18 Å in tRNA_open_ to accommodate the reoriented acceptor stem ***D,*** Along with the movement of the T-arm, the D-arm also rotates by 18° and shifts at the most distant point by 8 Å.

The tRNA_open_ and tRNA_closed_ structures are most clearly distinguished by the distinct orientation of their acceptor stem relative to the tRNA body and bound Trm10 (**Fig. 2B** and **Supplementary Fig. S9**). In tRNA_open_, the acceptor stem comprising base pairs G1-C70 to G7-C64 (along with the bases U57-U65 and U8 of the T-arm) is rotated by ∼23° and shifted at its maximum displacement by ∼18 Å from the corresponding position in tRNA_closed_. The movement of the acceptor stem is enabled by reorganization of T-arm nucleotides C46-U58 which act as a hinge region and are also rotated (41°) and shifted (18 Å) from their corresponding positions in tRNA_closed_ (**Fig. 2C**). The T-arm movement also results in a rotation (18°) and shift (8 Å) of the D-arm (G10-A23) to maintain the overall architecture of tRNA_open_ (**Fig. 2D**). The structures of the two monomeric Trm10-tRNA^Gly-GCC^ complexes thus reveal structurally distinct open and closed conformation of the substrate tRNA which arise from coordinated reorganizations of the acceptor stem, T-arm, and D-arm. Collectively, these observations indicate that Trm10 can induce distinct tRNA conformations that may contribute to the mechanism of substrate tRNA recognition and G9 modification.

### Contacts made by multiple conserved Trm10 residues direct tRNA binding

Trm10 binds tRNA^Gly-GCC^ using a positively charged surface with an opening to the catalytic site of the enzyme positioned to accommodate G9 for modification (**Fig. 3A-C**). Several Trm10 residues contact the tRNA to precisely position the substrate (**Fig. 3D-G** and **Supplementary Figs. S4** and **S10**). R121 forms a cation-π interaction with G10 (**Fig. 3D,E**), directly stabilizing the tRNA backbone conformation adjacent to the target nucleotide G9, which is flipped into the active site. Residues R127 (highly conserved in fungi) and R128 (found in some but not all Trm10 orthologs; **Supplementary Fig. S4**) also interact with the tRNA phosphate backbone of A27/A28 and C25/C26/A27 in the anticodon stem near the tRNA core (**Fig. 3D,E,H** and **Supplementary Fig. S4** and **Table S1**). Substitution of these residues with glutamic acid in *S. cerevisiae* Trm10 results in loss of methylation activity (38). Our structures thus reveal the molecular basis for the important contributions of these residues in stabilizing the distorted tRNA structure surrounding the target site in *S. cerevisiae* Trm10 (38,51).

**Fig. 3.**
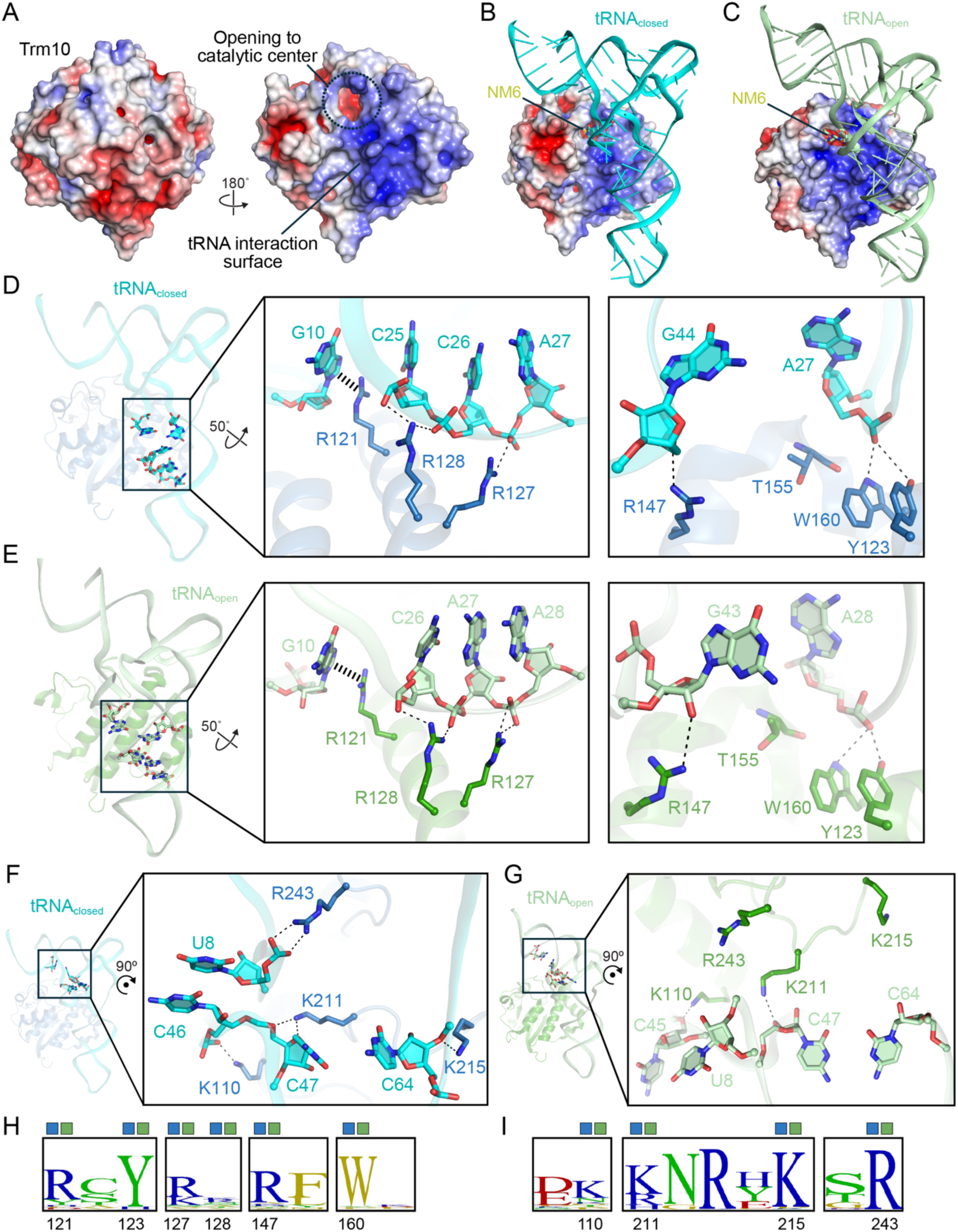
Trm10 engages with tRNA via conserved basic residues. ***A-C,*** Trm10 possesses a positively charged surface that facilitates interaction with the tRNA and guides the flipped G9 into the active site opening. ***D,*** In tRNA_closed_, side chains of R127 and R128 contact the negatively charged phosphate group of A27 and C25/C26, respectively, to stabilize the anticodon stem of the tRNA. R121 contacts the base of G10 in the D-arm. A second group of residues−R147 and T155/W160/Y123−is positioned adjacent to the phosphate of G44 (part of the variable loop) and A27, respectively, also contributing to tRNA stabilization. These interactions described in the main text are denoted with dashed lines. ***E,*** Same view as for panel D, but for the Trm10-tRNA_open_ complex showing that R127 and R128 interact with A28 and C26/A27, respectively. Similarly, R147 and T155/W160/Y123 remain in position to contact the RNA backbone but are adjusted to contact G43 and A28, respectively. Interactions in the acceptor stem region in the ***F***, Trm10-tRNA_closed_ and ***G***, Trm10-tRNA_open_ complexes show the side chains of amino acids K211 and K110 contacting C47 and C45 (tRNA_open_)/C46 (RNA_closed_), respectively. In tRNA_closed_, R243 and K215 also contact U8 and C64, respectively, which are missing in tRNA_open_ due to the movement of the acceptor stem. ***H,*** Sequence comparison of *S. cerevisiae* Trm10 and 321 Trm10 homologs from fungi depicts the conservation of R121, Y123, R127, R128, R147, and W160. Letter coloring denotes basic (blue), acidic (red), non-polar (green) and hydrophobic (gold) residues. The symbols above the alignment indicate (where present) that the residue sidechain position was directly supported by the map in the tRNA_open_ (green) and/or tRNA_closed_ (blue) complexes or inferred from the modeled backbone. ***I,*** As for panel H, but for residues K110, K211, K215, and R243.

Several other positively charged, polar, and aromatic residues form nearby interactions with the tRNA. R147 interacts with variable loop nucleotides via either the phosphate of G44 (tRNA_closed_) or the ribose 2’-OH of G43 (tRNA_open_). Additionally, W160 and Y123 interact with the phosphate of either A27 or A28 in tRNA_closed_ and tRNA_open_, respectively, and the T155 peptide backbone packs against the RNA in this region (**Fig. 3D,E**). Consistent with these interactions, Y123, R147, and W160 are highly conserved among fungal Trm10 homologs, while T155 exhibits very low conservation (**Fig. 3H** and **Supplementary Fig. S4** and **Table S1**). Notably, residues R121 and R147 appear to cooperatively engage the two strands of the tRNA from opposing sides, functioning like a “molecular pincer” to grip the tRNA. Together with the other adjacent residues, these contacts precisely position the substrate for G9 modification (**Supplementary Fig. S11A**).

At a second site of interactions with the tRNA adjacent to the acceptor stem and variable loop, the moderately conserved K110 interacts with either C46 (tRNA_closed_) or C45 (tRNA_open_) in the variable loop, while K211 is also positioned close to C47 as well as the phosphate of G9 in both tRNA conformations (**Figs. 3F,G,I** and **4**, and **Supplementary Fig. S4** and **Table S1**). K211 is thus positioned to stabilize the reconfigured tRNA backbone of the flipped target base in a conformation that appears to be further stabilized by interaction of R243 and the phosphate of the adjacent U8 in tRNA_closed_. Interestingly, the interaction with R243 is absent for tRNA_open_ as the acceptor stem moves away from Trm10 (**Fig. 3G**). Similarly, K215 forms a hydrogen bond with the phosphodiester backbone of C64 in tRNA_closed_ that is absent for tRNA_open_ due to the rotation of the T-arm (**Fig. 3F,G**).

Similar to R121 and R147, in tRNA_closed_ residues K110, K211, and R243 appear to act as molecular pincers to secure the tRNA (**Supplementary Fig. S11B**). However, in contrast to the moderate conservation of K110 and K211 in fungi, R243 is highly conserved in >95% of analyzed sequences (**Supplementary Fig. S4** and **Table S1**). Moreover, despite varied conservation of K211 across eukaryotes, an analogous K185 residue was also implicated in tRNA binding in studies of *S. acidocaldarius* Trm10 with tRNA, consistent with some conserved features of tRNA binding (17). Although tRNA substrate specificities of fungal Trm10 enzymes remain incompletely defined, variation in some of these residues may contribute to differences in tRNA isotype selection across species, a hallmark of m^1^G9 methylation (16). Nonetheless, electrostatic complementarity and residue conservation support a model in which Trm10 uses a set of basic and other residues to engage extensively with different regions of its tRNA substrate, with several acting in a coordinated fashion, to stabilize the distorted RNA conformation and position the substrate for accurate G9 modification.

### Target nucleotide G9 is flipped into the active site

To make the N1 target atom accessible for modification, the G9 base is flipped out of the tRNA core and positioned in the active site (**Fig. 4A** and **Supplementary Fig. S12**). The conserved and functionally critical Q118 (15) stabilizes G9 in its flipped conformation through interactions with the nucleobase N3 and primary amine group (exocyclic N2) (**Fig. 4B-D**, and **Supplementary Fig. S4** and **Table S1**). Additionally, S114, which is highly conserved in fungi, interacts with both the G9 ribose 2’-OH and the phosphodiester backbone. In TRMT10C, structurally analogous N222 makes a corresponding interaction with the G9 ribose (33), suggesting a partially conserved nature of this interaction among the human proteins (**Supplementary Fig. S13A**). Finally, the G9 exocyclic amino group interacts with either the OH group of T248 (tRNA_closed_) or T247 (tRNA_open_) (**Fig. 4C,D**). However, both Thr residues are only modestly conserved fungal Trm10 enzymes and are typically replaced by non-polar hydrophobic residues in other family members (**Fig. 4B** and **Supplementary Table S1**), suggesting that they may be important for enclosing the flipped base within the active site through shape complementarity rather than direct hydrogen bonding. Indeed, a notable feature in the active site of Trm10 is the presence of a highly conserved hydrophobic pocket, in which the base of G9 packs against I208, V209, and V245–Trm10 amino acids that are functionally conserved across fungal homologs (**Figs. 4B,E,F**, and **Supplementary Fig. S4** and **Table S1**). Additionally, in tRNA_open_, the backbone of I208 and V245 forms hydrogen bonds with G9 (**Fig. 4F**). The positions of the same residue sidechains were not directly visible in the in tRNA_closed_ map but were instead modeled based on the backbone placement; thus, it is unclear whether the interactions formed by these residues are specific to the tRNA_open_ conformation.

**Fig. 4.**
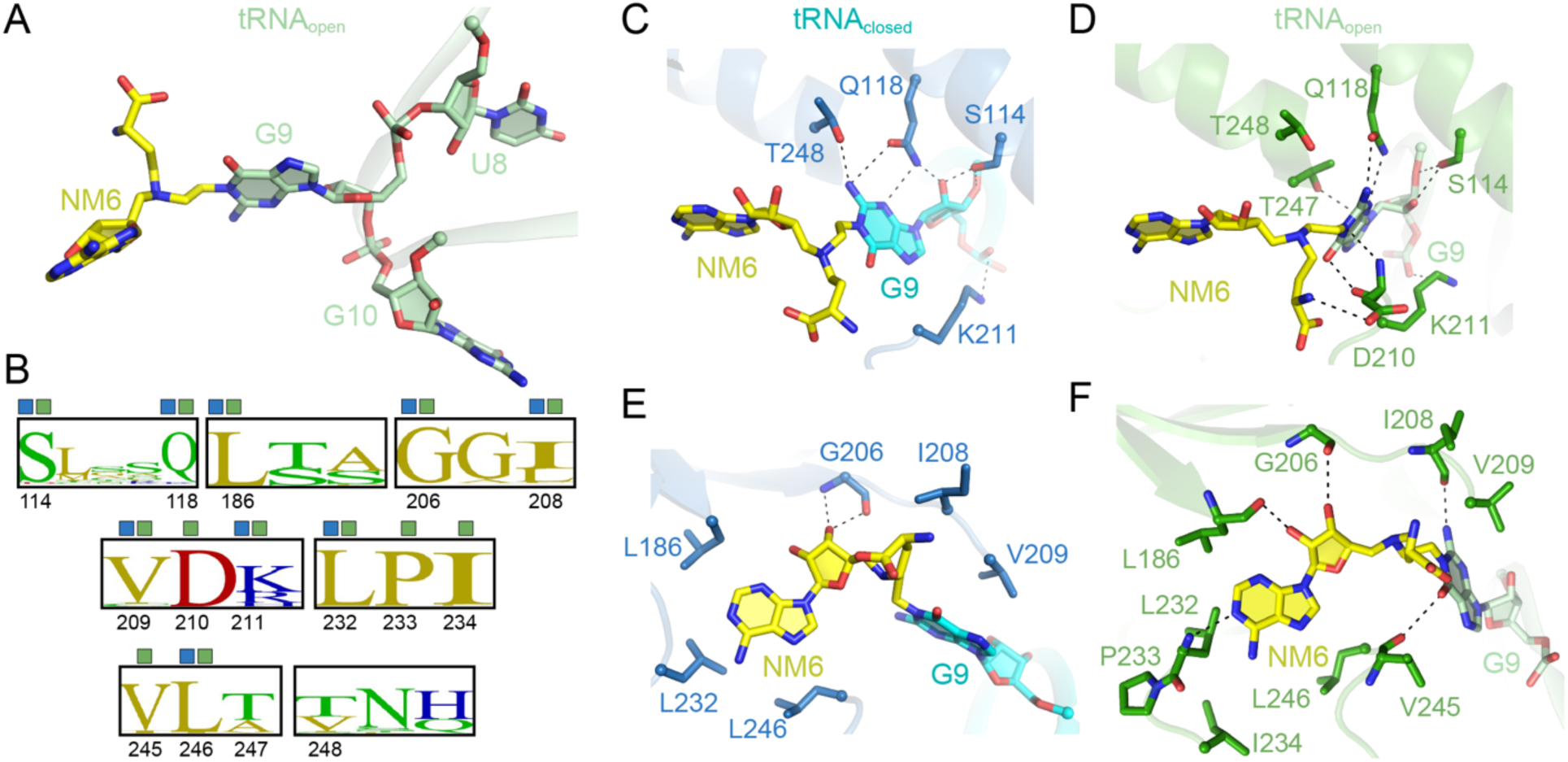
Conserved polar and hydrophobic residues in the active site stabilize flipped G9-NM6. ***A***, The G9 bound to NM6 adopts a flipped orientation compared to the adjacent nucleotides U8 and G10. ***B,*** Sequence alignment of *S. cerevisiae* Trm10 and fungal Trm10 homologs reveals strong conservation across species among key residues involved in stabilizing the flipped nucleotide. Letter coloring denotes basic (blue), acidic (red), non-polar (green) and hydrophobic (gold) residues. The symbols above the alignment indicate (where present) that the residue sidechain position was directly supported by the map in tRNA_open_ (green) and/ or tRNA_closed_ (blue) complexes or inferred from the modeled backbone. ***C-D***, In tRNA_closed_, side chains of polar residues, T248, Q118, S114, and K211 contact G9. In tRNA_open_, these contacts are maintained, but with T247 in place of T248, and an additional interaction is observed between D210 and both G9 and NM6. ***E-F,*** G9 and NM6 establish base stacking interactions with primarily non-polar residues, as well as direct contacts with G206, L186, V245, and I208.

The covalent attachment formed between the SAM analog NM6 and the G9 nucleobase N1 position during catalysis is clearly observed in the map (**Fig. 1A,B**). NM6 adopts the typical bent conformation observed for SPOUT methyltransferase-bound cosubstrates (52), and aligns closely with SAH observed in the previously reported yeast Trm10 SPOUT structure (**Supplementary Fig. S13B,C**) (15). The bent conformation of NM6 is accommodated within a highly conserved hydrophobic pocket, which also forms part of the trefoil knot structure (53). In the tRNA_closed_ structure, side chain packing interactions are observed with L186, L232, and L246, and backbone interaction with G206 (**Fig. 4E**). The map for the Trm10-tRNA_open_ also allowed modeling of additional interactions between NM6 and the main chain atoms of L186 and L232, and positioning of the I234 side chain to enclose its adenine ring (**Fig. 4F**). Finally, the carboxylate side chain of D210 interacts with the methionyl amino group of NM6 in tRNA_open_, while the amino acid backbone atoms stabilize G9 (**Fig. 4D**). A similar interaction is observed for the modeled side chain in the tRNA_closed_ complex, although the corresponding density is absent. The selective interaction of this highly conserved amino acid with the amino acid portion of the cofactor further emphasizes an important role for this residue in positioning reactive groups for catalysis rather than a direct role in catalysis, in agreement with biochemical studies ruling out an essential role for this residue as a general base, as had been suggested initially (18,31,51).

These structures thus reveal how a collection of conserved polar and hydrophobic residues in Trm10 coordinate positioning of both the flipped G9 base and the SAM analog to enable efficient and accurate methylation by Trm10.

### Q118 mediates selective stabilization of G9 for methylation

To better understand how Trm10 selectively recognizes and modifies G9, we compared a portion of the active site of TRMT10C in complex with tRNA^Ile^ (PDB code 8CBO; contains G9) or tRNA^His,Ser^ (PDB code 8CBK; contains A9) (34), and the Trm10-tRNA_open_ complex in which interactions with G9 and NM6 were best defined by our maps. Additionally, to enable these comparisons, NM6 was removed and replaced by SAM (Trm10-tRNA_open-G9_), and a second model was then generated in which G9 was replaced with A9 (Trm10-tRNA_open-A9_). For both TRMT10C and Trm10, the amino and carboxyl groups of the Q226/Q118 side chain contact G9 (**Fig. 5A**). A similar interaction is maintained by TRMT10C with A9 but is absent in the modeled Trm10-tRNA_A9_ complex due to the movement of the nucleobase away from Q118 (**Fig. 5B**). In TRMT10C, both G9 and A9 are contacted by N348 with additional interactions made by N350 or D314, respectively (**Fig. 5A,B**). The structurally corresponding Trm10 residues, R243, V245, and D210, are unable to make similar interactions due to the hydrophobic side chain of V245 and distinct orientations of D210 and R243, which place them beyond hydrogen bonding distance.

**Fig. 5.**
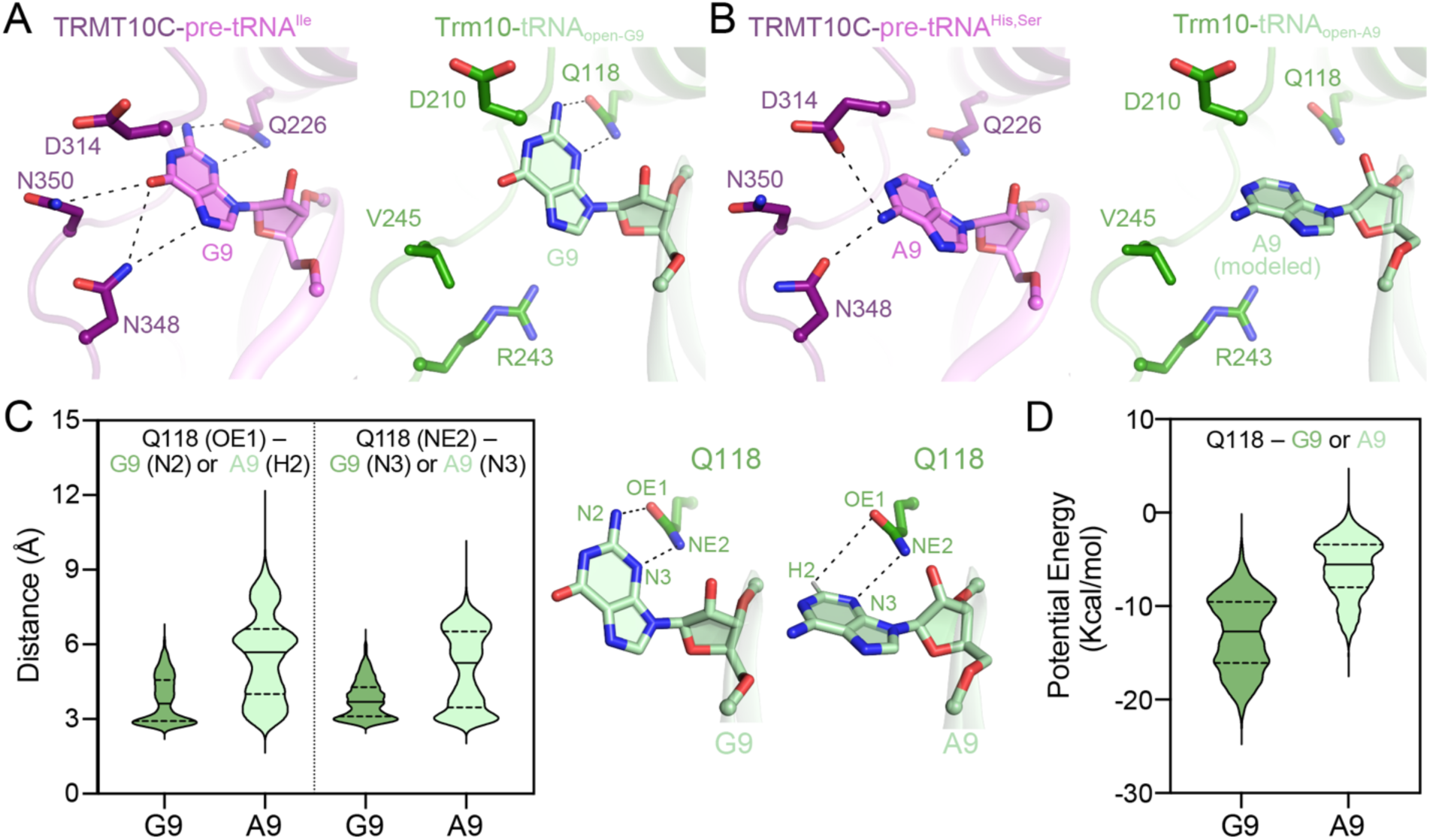
Q118 is responsible for guiding selective modification of G9 by Trm10. ***A,*** Active site comparison of TRMT10C-pre-tRNA^Ile^ (*left*, G9, violet, PDB 8CBO) and *S. cerevisiae* Trm10–tRNA^Gly-GCC^ (*right*, in the tRNA_open-G9_ model, green) showing the conserved contact between Q226/Q118 and G9. ***B***, This interaction is retained with A9 in the TRMT10C–pre-tRNA^His,Ser^ complex (*left*, A9, violet, PDB 8CBK) but absent in the modeled Trm10–tRNA^Gly-GCC^ complex (*right*, in the tRNA_open-A9_ model) due to nucleobase movement away from Q118. Additional TRMT10C residues involved in interaction are N348 (both G9 and A9), N350 (G9), D314 (A9), whereas the corresponding Trm10 residues (R243, V245, D210) fail to form equivalent hydrogen bonds due to unfavorable polarity and geometry. ***C,*** MD simulations of Trm10– tRNA_open-G9_ and Trm10–tRNA_open-A9_ complexes show that the two side chain interaction distances of Q118 with G9 are significantly shorter and within hydrogen bonding distances compared to that with A9. ***D,*** Potential energy profile of the Q118-G9 interaction is lower than that with A9, suggesting a more stable interaction with guanosine.

We next used MD simulations of the Trm10-tRNA_open-G9_ and Trm10-tRNA_open-A9_ complexes and monitored the distance distribution between the side chain of Q118 and G9/A9 over the course of each trajectory. This analysis reveals that the Q118-G9 interaction is significantly more stable, with a distribution centered on a shorter distance and retaining consistent hydrogen bonding or electrostatic interaction (**Fig. 5C**). In contrast, the Q118-A9 distance distribution is shifted toward longer distances, indicating a weaker or less persistent interaction over the simulation (**Fig. 5C**). This comparison thus suggests that Q118 engages more consistently in favorable interactions with G9 compared to A9. We also calculated the total potential energy profiles of each complex, with the G9-containing system displaying a lower average energy and reduced variation over the simulation (**Fig. 5D**). This result is again consistent with the presence of G9 resulting in a more stable conformational ensemble, likely due to the role of Q118 in anchoring the nucleotide in a position suitable for modification. In contrast, the lack of recognition of A9-containing tRNAs by *S. cerevisiae* Trm10 can be explained by the less favorable interaction with the nucleotide, along with a higher energy landscape.

Notably, this proposed role is also consistent with the larger effect on the observed rate of m^1^G9 modification (42-fold) *vs.* the rate of m^1^A9 modification (4-fold) upon alteration of the analogous residue Q122 (Q118 in *S. cerevisiae*) to alanine in the context of the bifunctional *T. kodakarensis* Trm10 (18).

Overall, these findings support a model in which Q118 selectively recognizes and stabilizes G9 for modification, a hallmark of fungal Trm10 enzyme specificity. Further, Trm10 enzymes from Archaea and eukaryotes with different specificities (A9 or dual G9/A9) also employ the analogous residue to Q118, but specificity appears to be supported by critical interactions made by other residues surrounding the target nucleotide. For example, we speculate that the selective recognition mechanism of A9 in human TRMT10B, the structure of which is unknown, may arise via interactions with Q148 (Q118 in *S. cerevisiae*) in combination with other residues in a manner similar to TRMT10C.

### A subpopulation of complexes containing two Trm10 enzymes is observed with a single tRNA

All RNA-modifying SPOUT family methyltransferases other than Trm10 have been shown to act as dimers (6,52). Interestingly, although the previously published yeast Trm10 enzyme-only structures (15) and our Trm10-tRNA complexes show a predominantly monomeric enzyme, a small subset of 2D classes in our dataset appeared to contain two Trm10 proteins for each tRNA molecule, i.e., (Trm10)_2_-tRNA^Gly-GCC^ (**Supplementary Fig. S2A**). The number of particles comprising these classes (∼5.2%) was significantly less than for monomeric Trm10, suggesting that a second Trm10 may only be present as a transient binding partner with the monomeric Trm10-tRNA complex. A 3D reconstruction of the (Trm10)_2_-tRNA^Gly-GCC^ complex revealed one Trm10 SPOUT domain bound to the tRNA core and positioned similarly to the enzyme in the monomeric Trm10-tRNA complexes, while the second Trm10 (Trm10’) is bound near the ASL but does not appear to contact the first Trm10 protomer (**Fig. 6A,B**), making this complex distinct from the typical SPOUT homodimer interaction. Although the map provided sufficient density to confidently dock and model the Trm10’ SPOUT domain, the quality of the overall map is affected by the significant orientation preference compared to the monomeric Trm10 complexes (**Supplementary Fig. S3G-I**), limiting our ability to definitively identify specific residues in tRNA or protein involved in the additional interaction.

**Fig. 6.**
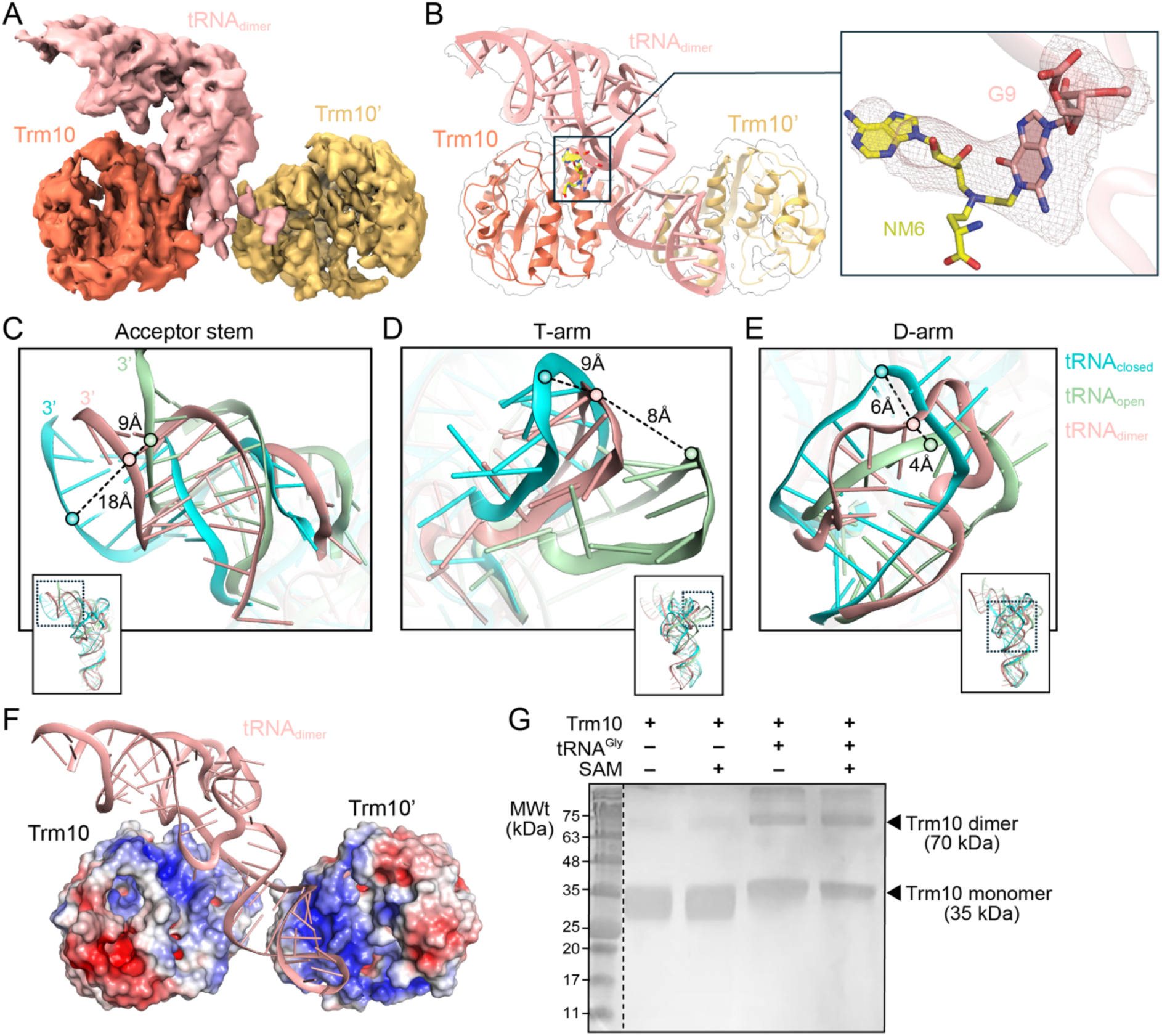
Trm10 dimer formation is facilitated by tRNA. ***A,*** CryoSPARC (CS)-sharpened map of (Trm10)_2_-tRNA complex (two Trm10s in orange and yellow; tRNA in light red) at 3.89 Å (threshold: 0.019; map value range: −0.0897 to 0.164). ***B,*** The final model is shown within a semi-transparent white map. ***Inset***, Close-up view of the NM6 (yellow) attached to G9 with the corresponding CS-sharpened map shown (threshold: 0.021). ***C-E,*** Comparison of the acceptor stem, D-arm, and T-arm of the tRNA_open_ (light green), tRNA_closed_ (cyan), and dimeric states indicate that the tRNA in those domains occupies an intermediate position compared to the corresponding regions in tRNA_open_ and tRNA_closed_. ***F,*** The two Trm10 protomers employ a positively charged surface to use the tRNA as a scaffold. ***G,*** Crosslinking assays using BS3 show that Trm10 remains monomeric in the absence of tRNA, but forms dimers only when tRNA is present.

The two Trm10s in the dimeric complex are structurally similar to the previously published Trm10 structure (**Supplementary Fig. S14A**), with an RMSD of ∼3.0 Å for both C_α_ atom alignments. In contrast, alignment of the tRNA bound to the two Trm10s to tRNA^Phe^ shows an RMSD of 7.8 Å over 63 phosphate atoms, with significant differences distributed across the acceptor stem, D-arm, and T-arm (**Supplementary Fig. S14B**). Comparing the tRNAs from the two monomeric complexes and the (Trm10)_2_-tRNA complex reveals the tRNA bound to two

Trm10 enzymes to be in an intermediate conformation in terms of the deformations observed in the acceptor stem, D-, and T-arm of tRNA_closed_ and tRNA_open_ (**Fig. 6C-E**). This observation suggests that the tRNA conformation in the (Trm10)_2_-tRNA complex may represent an intermediate stage in the process of recognition of tRNA by Trm10. Protein electrostatic surface calculation reveals that Trm10’ also employs a positively charged surface to interact with the tRNA, such that the tRNA is sandwiched between positive surfaces from the two proteins and further supporting the plausibility of the positioning of the second Trm10 molecule (**Fig. 6F**).

Finally, superimposing the AlphaFold full-length Trm10 structure on each individual Trm10 protomer is possible with no major clashes with the SPOUT domain of the other experimentally positioned protomer, suggesting that the two protein molecules can simultaneously bind to the tRNA without steric hindrance (**Supplementary Fig. S7C**). When both full-length models are aligned to the dimeric structure the two NTDs clash in the center of their main helix (**Supplementary Fig. S7C**, *center*), but the position of this region is likely to be dynamic, e.g. via movement at the flexible loop between the NTD and SPOUT domain, and our functional data indicate this region is not critical for tRNA binding or Trm10 activity (**Supplemental Fig. S6B,C**). In contrast, the functionally important NTD region most proximal to the SPOUT domain is positioned near the tRNA stem-loop in both modeled full-length protein. It is also notable that Trm10’ shows fewer direct contacts with the tRNA than Trm10 (i.e. the protomer common to both the dimeric and monomeric complexes), suggesting that its dynamic NTD might play a more prominent role in this additional interaction. However, like the NTD from the first Trm10, this region is unresolved in our map corresponding to this post-catalytic complex.

To further support the observation of two Trm10 enzymes binding to the tRNA and exclude the possibility that the observation is an artifact of the structure determination process, we performed a protein crosslinking analysis using the bifunctional reagent BS3 for Trm10 in solution in the presence and absence of both tRNA^Gly-GCC^ and SAM. Exclusively monomeric Trm10 was observed both with or without the cosubstrate in the absence of tRNA, whereas a prominent band corresponding to the presence of two Trm10s appeared in a tRNA-dependent manner, regardless of the presence of SAM (**Fig. 6G**). This result demonstrates the necessity of the tRNA substrate to promote the binding of two Trm10 molecules and can explain the absence of dimerization with a second Trm10 molecule in previous structural studies of the enzyme in which tRNA was absent (15). Interestingly, the tRNA-dependent crosslinking was also observed for the Trm10 NTD variants Trm10-8A and Trm10-Δ47, but not Trm10-Δ63, correlating dimer formation with the observed activities of these proteins (**Supplementary Fig. S6D**). The importance of the NTD region most proximal to the SPOUT domain is also consistent with the proximity of this region in each protomer with each other and the tRNA ASL.

Direct identification and experimental validation of key contacts made by Trm10’ to the tRNA is hindered by both the lower resolution of the map for the dimeric complex and the fact that six of the Trm10’ residues (R121, R126, R127, N153, N156, and N159; collectively, “6A”) potentially contacting the tRNA are common to both protomers. We therefore again turned to MD simulations as this approach allows *in silico* 6A substitution in both protomers at the same time (**Supplementary Fig. S15**), or in each individual protomer (**Supplementary Figs. S16** and **S17**) to dissect the potential contribution of these six residues to complex formation.

When both protomers have the 6A substitutions, Trm10’-6A rapidly disengages from the tRNA (within 10 ns of the MD production run; **Supplementary Fig. S15**, *right*) consistent with these contacts being important for Trm10’ interaction with tRNA. In contrast, with only Trm10 as the 6A variant (Trm10’ as wild-type), both protomers remain engaged with the tRNA with a modest increase in overall RMSD only for Trm10-6A, reflecting a lesser dependence of the Trm10 protomer-tRNA interaction on these residues compared to Trm10’ (**Supplementary Fig. S16A**). At the residue level, Trm10-6A exhibits differences in local dynamics (as assessed by backbone RMSF) both adjacent to the sites of substitution and more distant in the protein (**Fig. 7A** and **Supplementary Fig. S16B,** *left*). RMSF differences in this complex are also observed in the Trm10’ protomer (which has wild-type sequence) and the tRNA (**Fig. 7A** and **Supplementary Fig. S16B,** *right*). Notably, these changes are in regions involved in the protein binding-induced tRNA conformational reorganization in the acceptor stem and the D-arm as revealed by our structures (**Fig. 2**). When alternatively placed only in Trm10’, the 6A substitutions do not detectably alter the overall structure of either protomer (as assessed by chain RMSD; **Supplementary Figs. S17A**) but Trm10’-6A disengages from the tRNA later in the simulation. Local changes in residue dynamics are similarly fewer in this context though some changes in tRNA residues are observed again at sites in the tRNA core and ASL relevant to the distortions observed in our structures (**Fig. 7B** and **Supplementary Fig. S17B**).

**Fig. 7.**
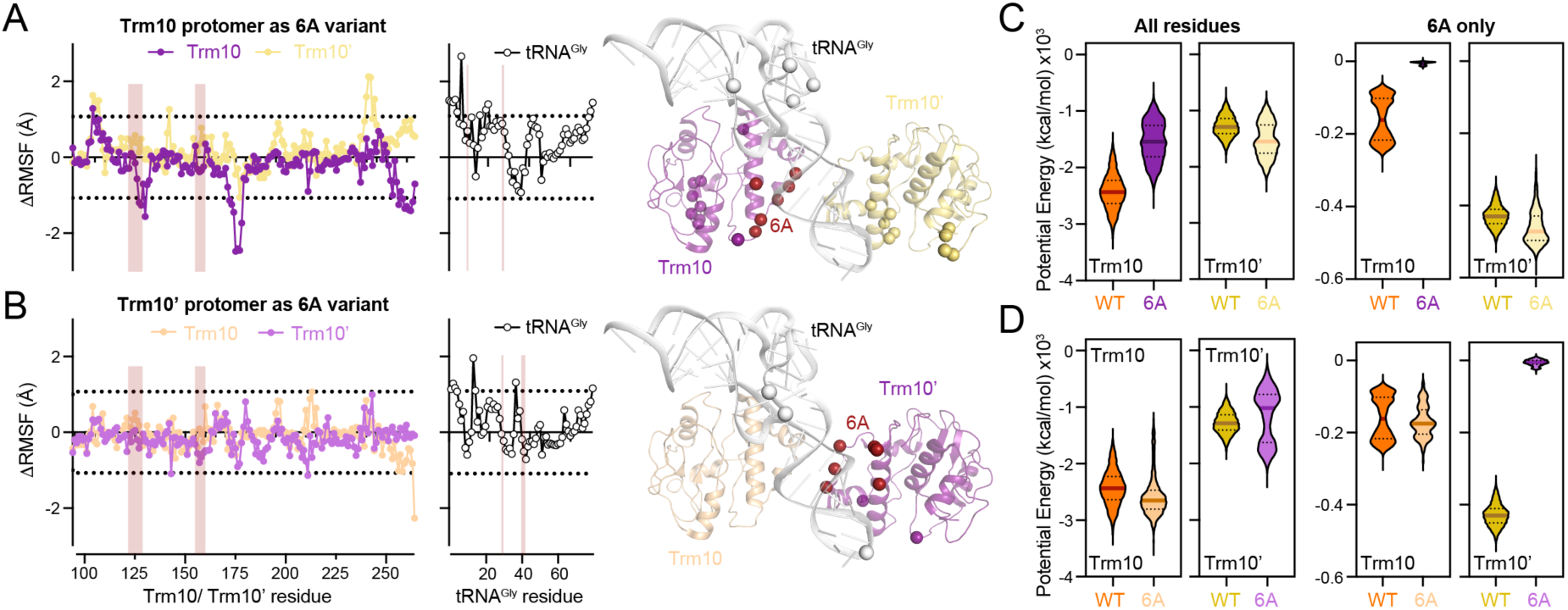
Six residues in the Trm10-tRNA interface stabilize the (Trm10)_2_-tRNA complex and may facilitate crosstalk between the protomers. ***A,*** Plots of difference RMSF (ΔRSMF) calculated by subtraction of RSMF values for the fully wild-type (WT) dimer from those of the 6A-containing dimer for the individual Trm10 (purple) and Trm10’ (yellow) protomer chains (*left*) and tRNA (*right*). Horizontal dotted lines indicate two standard deviations (up or down) from the average ΔRSMF. Residues outside these boundaries (corresponding to the greatest increases or decreases in residue dynamics) are shown on the structure as chain-colored spheres. Vertical shading on the plots indicates the locations of the amino acids in the protein chains and their sites of contact on tRNA. ***B,*** As for *panel A*, but with the 6A substitutions located in the Trm10’ protomer. ***C,*** Calculations of complex potential energy comparing the fully wild-type **and** 6A-containing dimer with the substitutions in the Trm10 protomer (purple) for all residues (*left two panels*) or the 6A residues only (*right two panels*). ***D,*** As for *panel C*, but with the 6A substitutions in Trm10’.

Finally, calculations of interaction energies, either for all residues or the 6A residues only, are also consistent with the importance of these contacts for each protomer engaging with tRNA. In each variant dimer context, the 6A substitutions result in a slight increase in the average calculated energies compared to wild-type when the changes are placed in either the Trm10 or Trm10’ protomer (**Fig. 7C,D**). Interestingly, however, there is also an associated decrease in energy in the non-mutated protomer of each variant dimer, further pointing to the potential for crosstalk between the two protomers of the dimeric complex which may control their coordinated action during tRNA modification.

In summary, while further high-resolution studies of the (Trm10)_2_-tRNA complex will be needed to identify unique Trm10’-tRNA contacts to enable experimental validation, our current structural, biochemical and computational studies support a model for Trm10 action that involves an unanticipated dimeric complex on tRNA. Further, our observations to date suggest that binding of the second Trm10 occurs only when tRNA is present, and that this recruitment affects the dynamics of both the tRNA and Trm10 protomer that modifies G9.

## DISCUSSION

Trm10 is an evolutionarily conserved tRNA methyltransferase that in yeast modifies the N1 base position of G9 in the tRNA core. In this work, we determined three distinct cryo-EM structures of the Trm10-tRNA complex: two monomeric complexes distinguished by the conformation of the bound tRNA, and a dimeric (Trm10)_2_-tRNA complex. The monomeric Trm10 complexes mark the first detailed structural insight into the process of tRNA substrate recognition and modification by this stand-alone monomeric SPOUT methyltransferase, captured on the tRNA in a state immediately after modification. Additionally, the unexpected dimeric (Trm10)_2_-tRNA complex revealed how two molecules of Trm10 may interact with a single tRNA. The structures allowed us to identify important interactions between Trm10 and tRNA that contribute to substrate recognition, stabilization of the flipped G9 nucleobase for modification, and insight into the mechanism of selective recognition of guanosine at position 9 by yeast Trm10.

The two Trm10 monomeric structures, sequence conservation analysis, and previous studies allow us to rationalize functional data from yeast, archaeal, and human proteins. For instance, the highly conserved D210/ D314/ D184 in the active site of *S. cerevisiae* Trm10, *S. acidocaldarius* Trm10, and human TRMT10C, respectively, are functionally important in all three enzymes (17,22,34,51). However, the lack of a direct interaction between the side chain of analogous D210 and the N1 atom of G9 in any of our structures is consistent with and rationalizes the previous demonstration that this residue does not function as a general base during catalysis in any stand-alone Trm10 enzyme from Eukarya or Archaea (31,51).

Furthermore, T247 interacts with G9 in its flipped conformation and alteration of the equivalent residue (T244) in *S. pombe* (15) reduced enzyme activity by 65%, underscoring the importance of properly positioning G9 for modification (**Fig. 4D**). Similarly, the V206A variant of *S. pombe* Trm10 (equivalent to V209 in *S. cerevisiae*) shows an 80% reduction in activity (15), indicating a key contribution of the G9 base stacking interaction for enabling target nucleotide modification (**Fig. 4E,F**).

Previous biochemical studies on residues implicated in tRNA binding showed that a double K153/R147E substitution affects tRNA binding in *S. pombe* (15) and our structures reveal R147 interacting with G43/G44, which would be disrupted by this substitution (**Fig. 3D,E**).

Additionally, substitution of K208 in *S. pombe* or K185 in S. *acidocaldarius* (equivalent to K211 in *S. cerevisiae*) to alanine or glutamate results in a ∼70% loss in activity compared to the wild-type enzyme (15,17), supporting a role for K211 in Trm10 activity via stabilization of the flipped G9 and interaction with C47 (**Figs. 3F,G** and **4C,D**, and **Supplementary Fig. S18B**). Interestingly, K211 of the second protomer (Trm10’) in the dimeric complex is also positioned to contact the tRNA such that this residue may play a dual role in tRNA substrate recognition and modification.

Similarly, a K121E substitution in *S. acidocaldarius*, equivalent to *S. cerevisiae* Trm10 R128 which contacts tRNA in both Trm10 protomers of the dimeric complex (**Supplementary Fig. S18**), results in impaired tRNA binding (17). Previous studies have shown that deletion of residues K236-R240 in *S. pombe* nearly abolished Trm10 activity (15). Notably, R243 in *S. cerevisiae*, which is equivalent to R240, interacts with U8 in the tRNA_closed_ conformation and thus appears to be involved in stabilization of tRNA architecture around the modified G9 base (**Fig. 3F**).

Our structures can also rationalize the underlying basis of disease-linked changes. Q118 is highly conserved in the active site of Trm10 and is important for enzymatic activity in *S. pombe* (15) and humans (TRMT10C) (54), and substrate selectivity in *T. kodakarensis*–observations reinforced and explained by our structures and MD simulations (**Figs. 4C,D** and **5**). A Q124K (Q118 in yeast) substitution in human TRMT10A has been reported in patients, but no associated physiological disorders have been observed (55). At G206, a residue important in stabilizing the bent cosubstrate conformation, clinical studies show that patients with mutations encoding an Arg or Ala substitution in human TRMT10A exhibit microcephaly and intellectual disability (24,56). Likewise, a nonsense mutation that results in truncation of TRMT10A at P233, another residue involved in positioning the cosubstrate, also results in neurological disorders (57). Trm10 residues K110, R121, and R127 are conserved in both *S. cerevisiae* and *S. pombe* and their substitution with glutamate was observed to affect either tRNA binding (*S. pombe*) (15) or methyltransferase activity (*S. cerevisiae*) (38), indicating the importance of these contacts. Additionally, clinical studies have shown substitutions of TRMT10A K116 (equivalent to K110 in *S. cerevisiae*) (58), R127 (equivalent to R121 in *S. cerevisiae*) (23), and R133 (equivalent to R127 in *S. cerevisiae*) (27) to be associated with microcephaly, intellectual disability, and epilepsy.

It is worth noting that most of the tRNA interactions identified in our structures are observed with the backbone (i.e., ribose and phosphate) moieties and are thus consistent with the lack of specific sequence-based identity elements for distinguishing substrate and non-substrate RNA, as has been demonstrated repeatedly for Trm10 enzymes (16,18,21). Rather, previous SHAPE analysis has shown that substrate recognition is dependent on inherent tRNA flexibility and the ability of Trm10 to induce distinct conformational changes in the tRNA upon binding (38). For instance, the D-arm was shown to be highly reactive in SHAPE studies which is reflected in our modevector analysis (**Fig. 2A**). Additionally, our studies indicated an unwinding of the helix in the ASL and a translational shift in the acceptor stem, while SHAPE showed an overall stabilization of both the regions. This suggests that Trm10 binding induces a conformational change that results in a more constrained ASL structure, possibly through interaction with the NTD as observed in our AF model, with reduced nucleotide flexibility and solvent accessibility in SHAPE, even though global backbone geometry is altered (**Supplementary Fig. S8**). Similarly, the lack of flexibility in the acceptor stem indicates a sterically constrained movement of the tRNA while sampling the different conformations observed in the tRNA_open_ and tRNA_closed_ forms. The relatively weaker patterns of conservation observed for Trm10 residues that interact with tRNA in the current structure may reflect the use of evolved changes in these amino acids to achieve the different patterns of tRNA isotype specificity across Trm10 from diverse organisms (**Supplementary Table S1**).

Interestingly, as more diverse orthologs of eukaryotic Trm10 enzymes are considered, the relative conservation of many of these residues also decreases. Human TRMT10A, the functional ortholog of *S. cerevisiae* Trm10, exhibits the most similarity to fungal Trm10s among the amino acids implicated above in tRNA interactions, except for K211, which is His in human TRMT10A (**Supplementary Table S1**). Perhaps not surprisingly, human TRMT10C lacks obvious conservation at several of these residues, which likely reflects its distinct composition as part of a multi-protein complex during its interaction with mitochondrial tRNA. Human TRMT10B represents somewhat of an intermediate among vertebrate enzymes, with some amino acids sharing similarity with fungal Trm10 and human TRMT10A, while others show distinct patterns of conservation. Therefore, the distinct substrate specificity for a single tRNA exhibited by this ortholog remains unexplained. Overall, the partial functional conservation of the tRNA-interacting residues suggests many of them are collectively important for tRNA binding. The species-specific absence of some individual Trm10-tRNA contacts is likely compensated by interactions of other residues or may reflect differences in how tRNAs are targeted for modification and could provide a compelling explanation for this so far unresolved aspect of Trm10 substrate selectivity.

Structural alterations in tRNA domains due to enzyme binding have been reported in studies of several other tRNA-modifying enzyme-substrate complexes (59–62), and particularly in tRNA methyltransferases, such as Trm5 (63), TrmD (64), and Trm6-Trm61 (65). While Trm5 and TrmD binding primarily affect the tRNA structure at the ASL, Trm6-Trm61, aspartyl-tRNA synthetase (AspRS) (60), and Sep-tRNA:Sec-tRNA synthase (SepSecS) (62) induce large distortions in the acceptor stem and T-arm regions. However, in contrast to the opening of the tRNA acceptor stem in the Trm10-tRNA_open_ complex, binding of AspRS, SepSecS, and Trm6-Trm61 induces a closure of the acceptor stem towards the tRNA body, representing an ‘induced fit’ mechanism of positioning the target nucleotide in the catalytic pocket for modification. Thus, while all these enzymes induce structural rearrangements surrounding their site of modification to facilitate catalysis, the conformational changes induced by Trm10 in the post-catalytic state are more distant from its modification site in the tRNA core.

The substantial tRNA distortions observed here may also reflect a Trm10-tRNA complex in a state poised for product release, a process that may require overall structural relaxation in the complex. In other systems, the free energy of an enzyme-substrate complex is lower in the catalytically active phase compared to the inactive phase or where the complex is not fully formed for catalysis (66,67). Accordingly, the Trm10-tRNA_closed_ complex exhibits lower calculated interaction energy compared to the tRNA_open_ form, and a more extensive buried surface (2330 *vs* 2080 Å^2^ in the complexes with tRNA_closed_ and tRNA_open_, respectively). Similar to the finding that the conformational change in dihydrofolate reductase during product release results in an increase in the free energy (68), our observations suggest that the tRNA_open_ may reflect a post-catalysis product release conformation. Similarly, a previously reported inactive AspRS-tRNA complex formed due to the absence of acceptor stem interaction with the enzyme (60), parallels our open Trm10-tRNA conformation and suggests functional inactivation before product (tRNA) release. Based on these observations and our three Trm10-tRNA structures, we propose an overall mechanism of catalysis and tRNA release, in which a monomeric Trm10 first binds the tRNA in a closed form for G9 methylation (**Fig. 8**), which is also similar to that seen in the TRMT10C-tRNA complex (34). During or following catalysis, the modified tRNA transitions to an open conformation, resulting in the release of the substrate from Trm10; however, post-catalytic dissociation of the complex is blocked in the current structure by the covalent attachment of NM6 to the modified tRNA. This model also offers a plausible explanation for why Trm10 binds to, but does not modify, the type II tRNAs. The structural transition observed in tRNA_open_ would be hindered by the extended variable loop in type II tRNAs, preventing the movement of the T-arm, and thereby restricting the tRNA to a catalytically unproductive closed conformation as suggested previously (16).

**Fig. 8.**
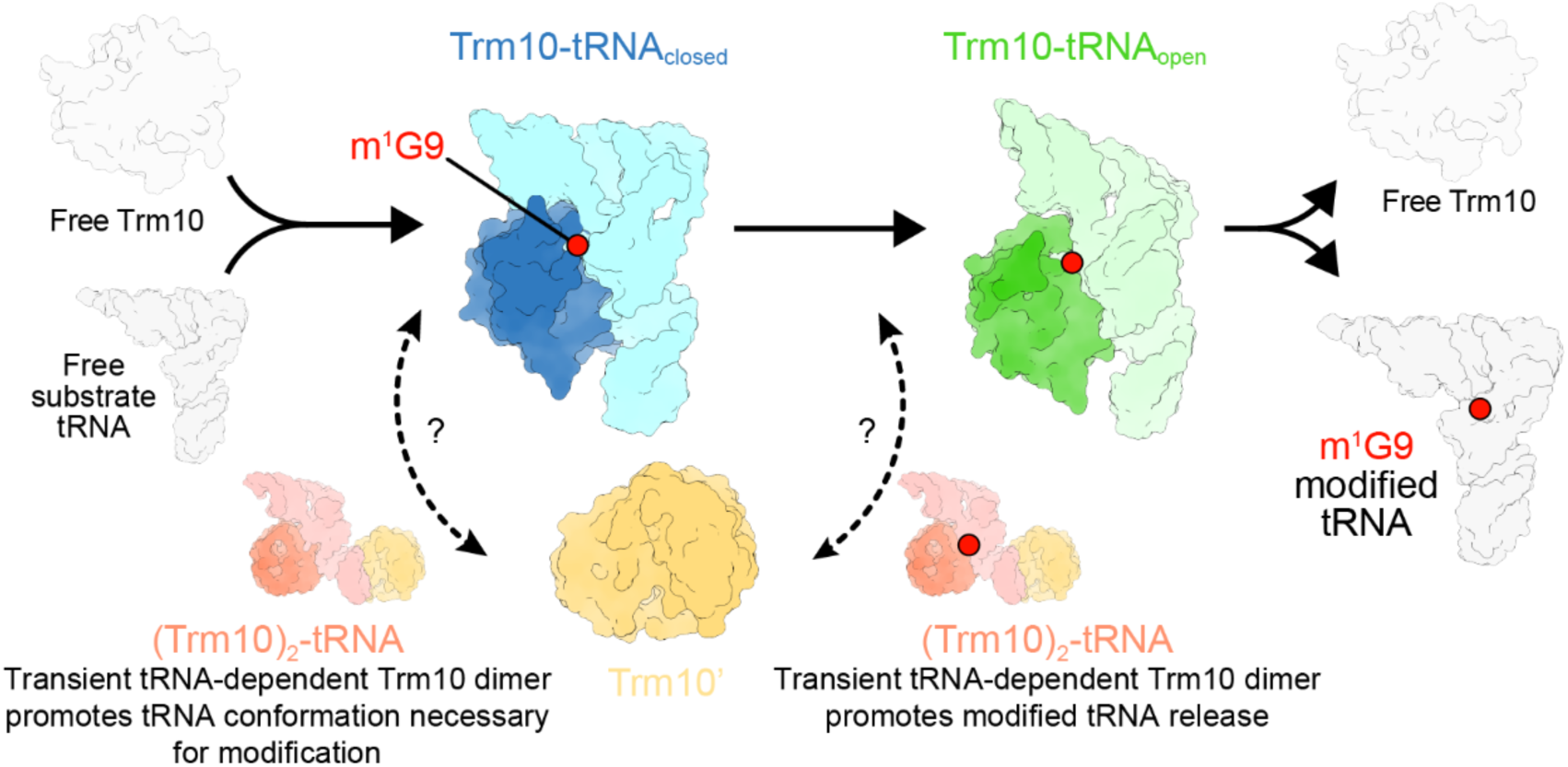
Proposed model for Trm10 catalysis mechanism and tRNA release. Free Trm10 (gray) and unmodified (G9) tRNA (gray) assemble in Trm10-tRNA_closed_ conformation (blue), promoting incorporation of the m^1^G9 modification (red sphere). Following catalysis, structural reorganization of the tRNA to the Trm10-tRNA_open_ conformation (green) enables release of free Trm10 and the m^1^G9-modified tRNA. Additionally, the transient (Trm10)_2_-tRNA complex may either promote the formation of a modification-competent tRNA conformation in the closed complex (*left*) or, following catalysis, promote opening of the tRNA structure to allow product release (*right*).

Finally, we also speculate that the (Trm10)_2_-tRNA complex may represent a transient functional state that either precedes the closed state by helping to position G9 for modification, or after modification, with the second Trm10 protomer binding the closed complex to displace the acceptor arm and weaken interactions to promote substrate release (steps indicted by dashed-line arrows with question marks in **Fig. 8**). Our speculation that the Trm10 second protomer may play a role in orienting G9 is supported by the structures of Trm6-Trm61 and Trm7-Trm734 bound to tRNA, in which Trm6 and Trm734 act as a tRNA-binding scaffold, while Trm61 and Trm7 provide the SAM binding motif and position the nucleotide to be modified (65,69). Consistent with this model, our MD studies of the (Trm10)_2_-tRNA complex highlight the influence of the Trm10’ protomer on the tRNA nucleotides involved in the tRNA_closed_ to tRNA_open_ structural transition. Together with the observed potential for communication between the two protomers, these findings suggest that the second Trm10 molecule plays a supportive role in guiding the formation or resolution of a catalytically competent Trm10–tRNA complex. However, further detailed studies will be necessary to fully define the molecular basis of tRNA substrate-dependent Trm10 dimerization, including protein-protein interactions, as well as the contribution of this unanticipated dimeric complex to the G9 modification process.

In conclusion, our studies reveal structures of the highly conserved atypical SPOUT methyltransferase Trm10 bound to a substrate tRNA in the absence of other binding partners and offer the first evidence of Trm10 activity employing a unique, transient tRNA-dependent dimeric complex to modify its substrate. Additionally, these studies shed light on the unique mechanism of selective recognition of G9 for modification, offering a mechanistic blueprint for substrate recognition by this atypical fungal tRNA SPOUT methyltransferase.

## DATA AVAILABILITY

Structural coordinates and EM maps have been deposited in the Protein Data Bank (PDB) and Electron Microscopy Data Bank (EMDB), respectively, with accession codes 9XZQ and EMD-72368 (Trm10-tRNA_closed_), 9XZR and EMD-72369 (Trm10-tRNA_open_), and 9XZS and EMD-72370 ((Trm10)_2_-tRNA). All other data are available in the main text or the supplementary materials.

## SUPPLEMENTARY DATA

Supplementary Data are available at online.

## ACKNOWLEDGMENTS

We thank Drs. Ed Eng, Eugene Chua, Aaron Owji, and Misha Kopylov from the New York Structural Biology Center (NYSBC) for their help with cryo-EM sample preparation and data processing. Some of this work was performed at NCCAT and the Simons Electron Microscopy Center located at the NYSBC, supported by the NIH Common Fund Transformative High Resolution Cryo-Electron Microscopy program (U24 GM129539), and by grants from the Simons Foundation (SF349247) and NY State Assembly. We also thank Drs. Srihari Koripella and Ricardo Guerrero-Ferreira from the Robert P. Apkarian Integrated Electron Microscopy Core of Emory University for their guidance with sample preparation and use of the facility’s electron microscopes.

