## Supplementary Materials for "Molecular basis of tRNA substrate recognition and modification by the atypical SPOUT methyltransferase Trm10"

**This PDF file includes:**

Supplementary Figures S1-S18
Supplementary Table S1

### SUPPLEMENTARY FIGURES

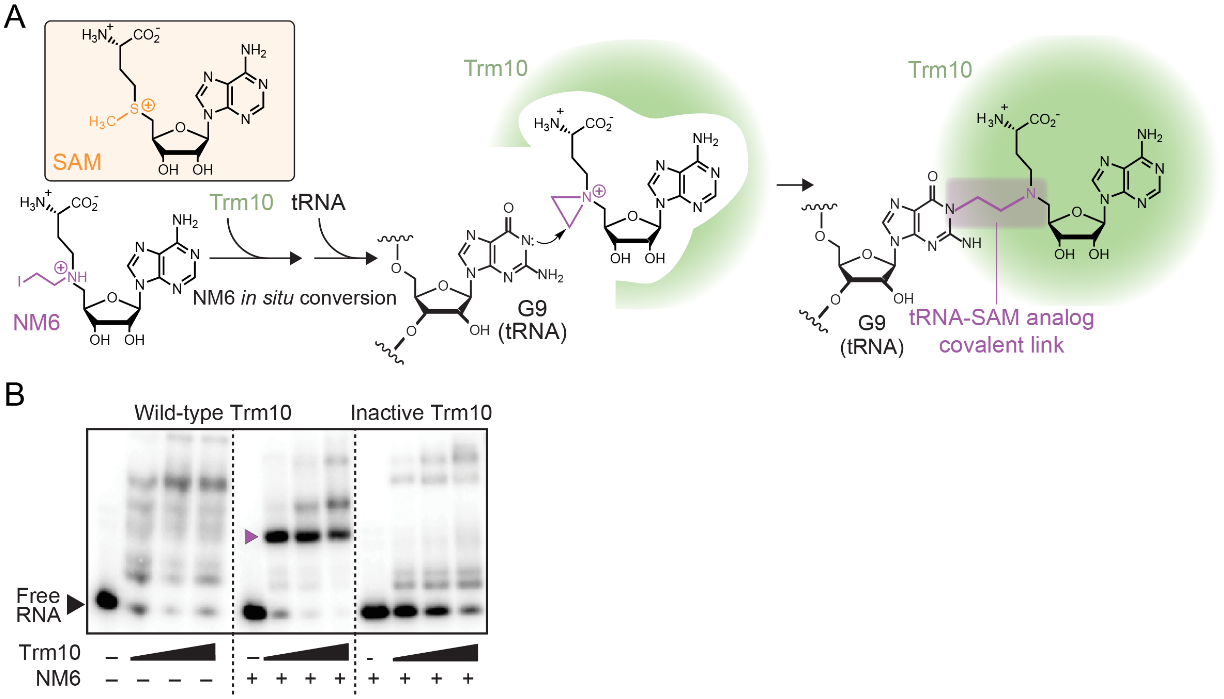


**Fig. S1. Covalent attachment of the SAM analog NM6 to tRNA captures the Trm10-tRNA complex in an immediately post-catalytic state. *A,*** Trm10 uses the S-adenosyl-L-methionine (SAM) analog "N-mustard 6" (NM6; *lower left*) as an alternative cosubstrate for Trm10-mediated tRNA modification. The analog is covalently attached to the N1 position of G9 during the catalytic reaction, trapping the Trm10-tRNA complex in a post-catalytic state due to the inherent affinity of Trm10 for the tRNA as well as the covalently attached NM6. ***B,*** Native PAGE analysis showing formation of a stable Trm10-tRNA complex only in the presence of NM6 (compare *left* and *center* panels; complex indicated by purple arrow). This complex is also absent without Trm10 catalytic activity to transfer NM6 to the tRNA (*right* panel; "Inactive Trm10" is the previously reported KRR variant-see Strassler *et al. J. Biol. Chem*. 2023 (Ref. 38)).

*[Figure is shown on the following page]*

**Fig. S2. Workflow for cryo-EM structure determination of the Trm10-tRNA$_{open}$, Trm10-tRNA$_{closed}$, and (Trm10)$_2$-tRNA complexes**. ***A,*** Sample micrograph, CTF estimation, particle picking, and example 2D classes. ***B***, 3D refinement workflow resulting in the final Trm10-tRNA complexes for tRNA$_{open}$ and tRNA$_{closed}$. *Ab-initio* refinements of the 2D classes produced 3D maps from which junk particles were removed by heterogeneous refinement. The best maps were used for non-uniform (NU) refinement and further polished by CTF refinement, reference-based motion correction, and DEM-sharpening, which produced a 3.37 Å map for the Trm10-tRNA$_{open}$ complex. For the tRNA$_{closed}$ complex, additional rounds of 3D classification and NU refinement yielded a 3.63 Å map. ***C,*** For the (Trm10)$_2$-tRNA complex, the best classes obtained from *ab-initio* refinement were further processed by NU refinement and reference-based motion correction, followed by multiple rounds of 3D classification to remove bad classes. Additional NU refinement and DEM sharpening produced a map of 3.89 Å.

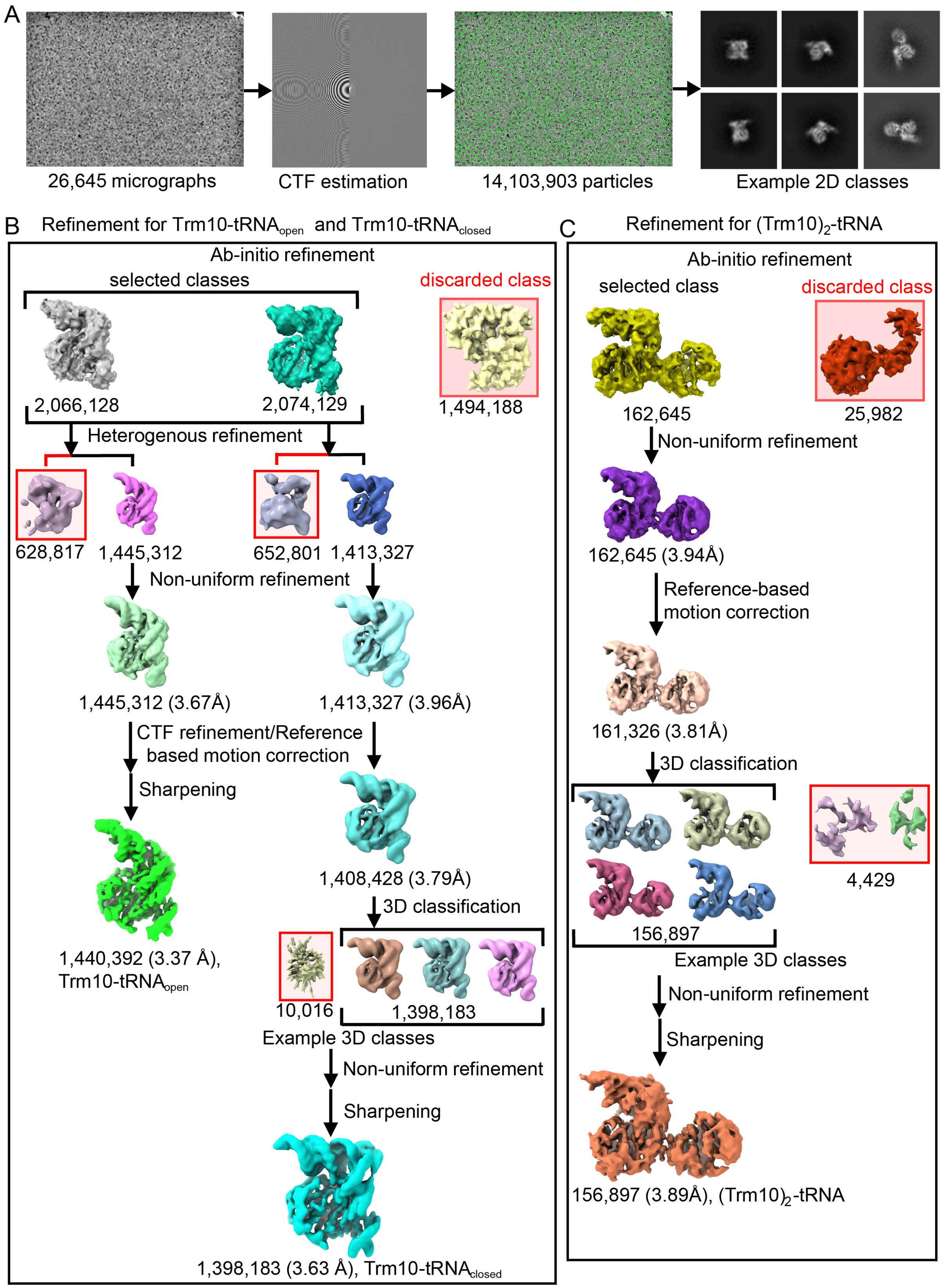


**Fig. S2. Workflow for cryo-EM structure determination of the Trm10-tRNA$_{open}$, Trm10-tRNA$_{closed}$, and (Trm10)$_2$-tRNA complexes**. *[Full legend is provided on the previous page]*

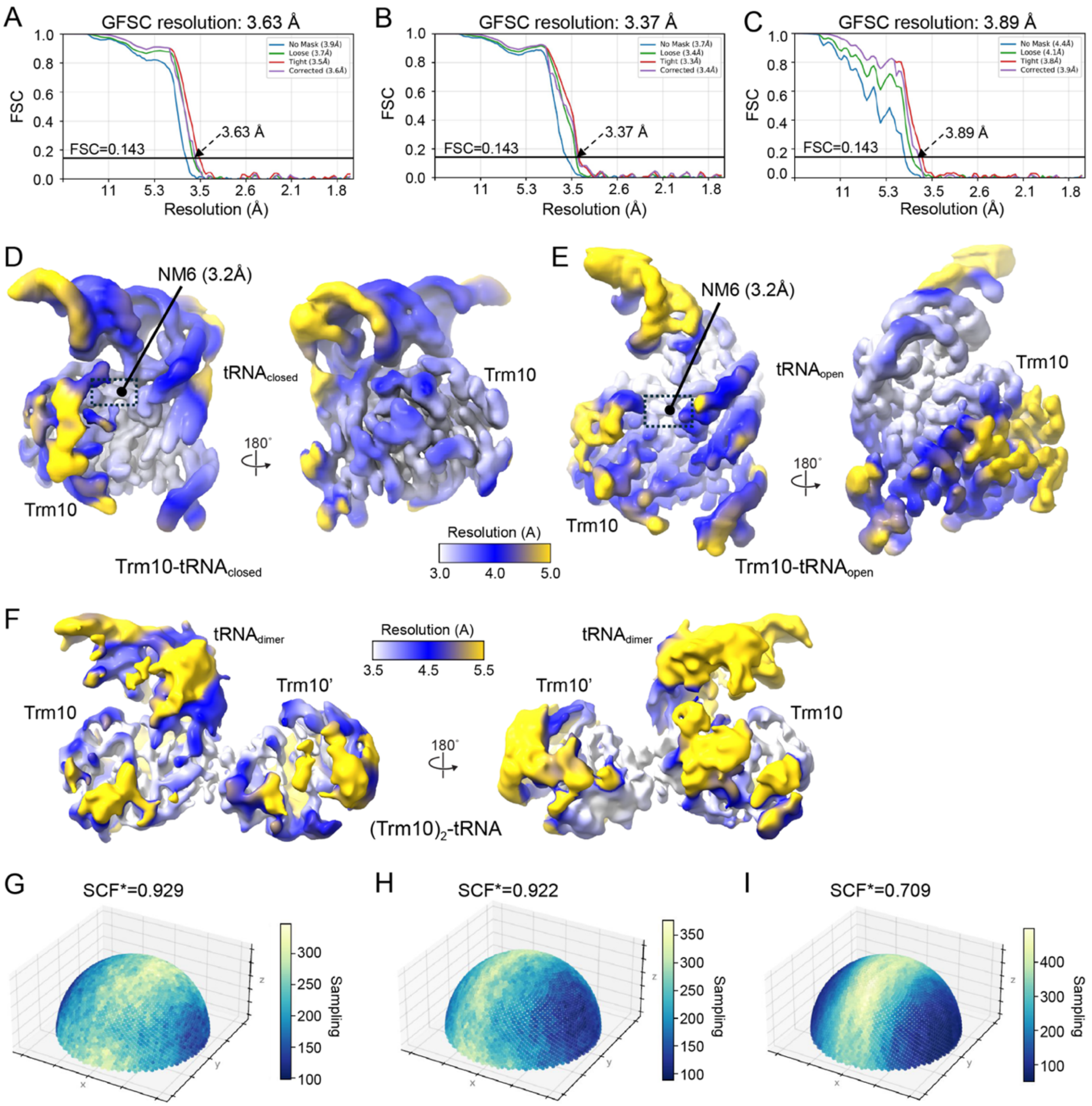


**Fig. S3. Analysis of cryo-EM map resolution.** Gold-standard Fourier Shell Correlation (GFSC) curves for the ***A,*** Trm10-tRNA$_{closed}$, ***B,*** Trm10-tRNA$_{open}$, and ***C,*** (Trm10)$_2$-tRNA complexes. The resolution of each map was determined from the 0.143 FSC intercept of the corrected data. Curves shown are for the unmasked map (blue), loosely masked map (green), tightly masked map (red), and corrected map (red). Differentially colored maps showing local resolution (values are indicated in the scale bar) of the ***D,*** Trm10-tRNA$_{closed}$, ***E,*** Trm10-tRNA$_{open}$, and ***F,*** (Trm10)$_2$-tRNA complexes. Sampling Compensation Factor (SCF) for the ***G,*** Trm10-tRNA$_{closed}$, ***H,*** Trm10-tRNA$_{open}$, and ***I,*** (Trm10)$_2$-tRNA complexes, showing significant orientation preference in the dimeric complex.

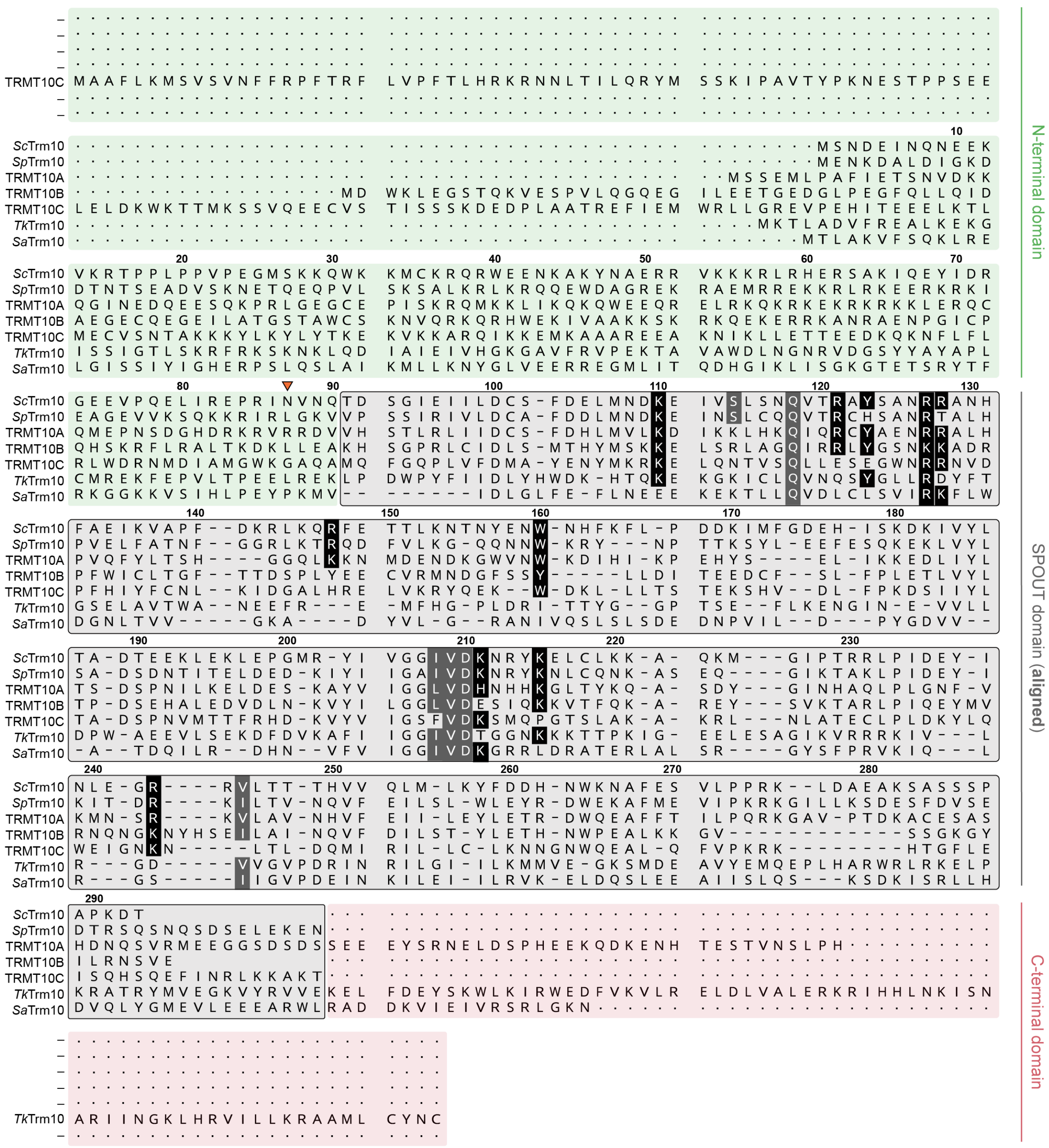


**Fig. S4. Protein sequence alignment for representative yeast, human and archaeal Trm10 enzymes.** Proteins were aligned on their conserved SPOUT domains (gray shading) and the non-conserved N- (green shading) and C-terminal (red shading) sequences were appended to the alignment. *S. cerevisiae* Trm10 (*Sc*Trm10) is shown in the top row with corresponding amino acid residue numbering; other enzymes are those from *Schizosaccharomyces pombe* (*Sp*Trm10), *Thermococcus kodakarensis* (*Tk*Trm10), *Sulfolobus acidocaldarius* (*Sa*Trm10), and human (TRMT10A, TRMT10B, and TRMT10C). Fully and partially conserved residues discussed in the main text that interact with tRNA or G9 specifically are highlighted on the alignment with white text and black and gray background shading, respectively; the first amino acid resolved in the map for the Trm10-tRNA complexes (N87) is indicated with an orange arrowhead.

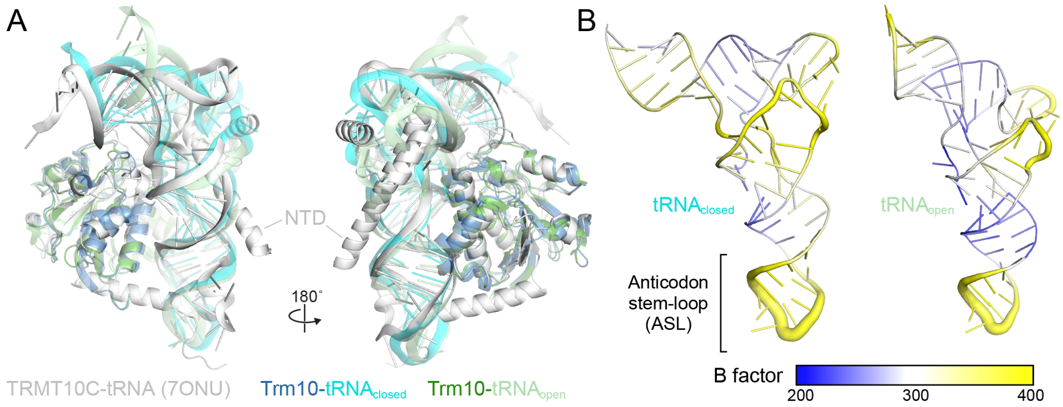


**Fig. S5. Structural features of the monomeric Trm10-tRNA$_{closed}$ and Trm10-tRNA$_{open}$ complexes.** ***A,*** In both complexes, *S. cerevisiae* Trm10 SPOUT (green and blue) binds to the tRNA (light green and cyan) in a manner closely resembling hTRMT10C (white), which also includes the helical NTD. ***B,*** Both the tRNA$_{open}$ and tRNA$_{closed}$ conformations exhibit elevated B factors in the anticodon stem loop.

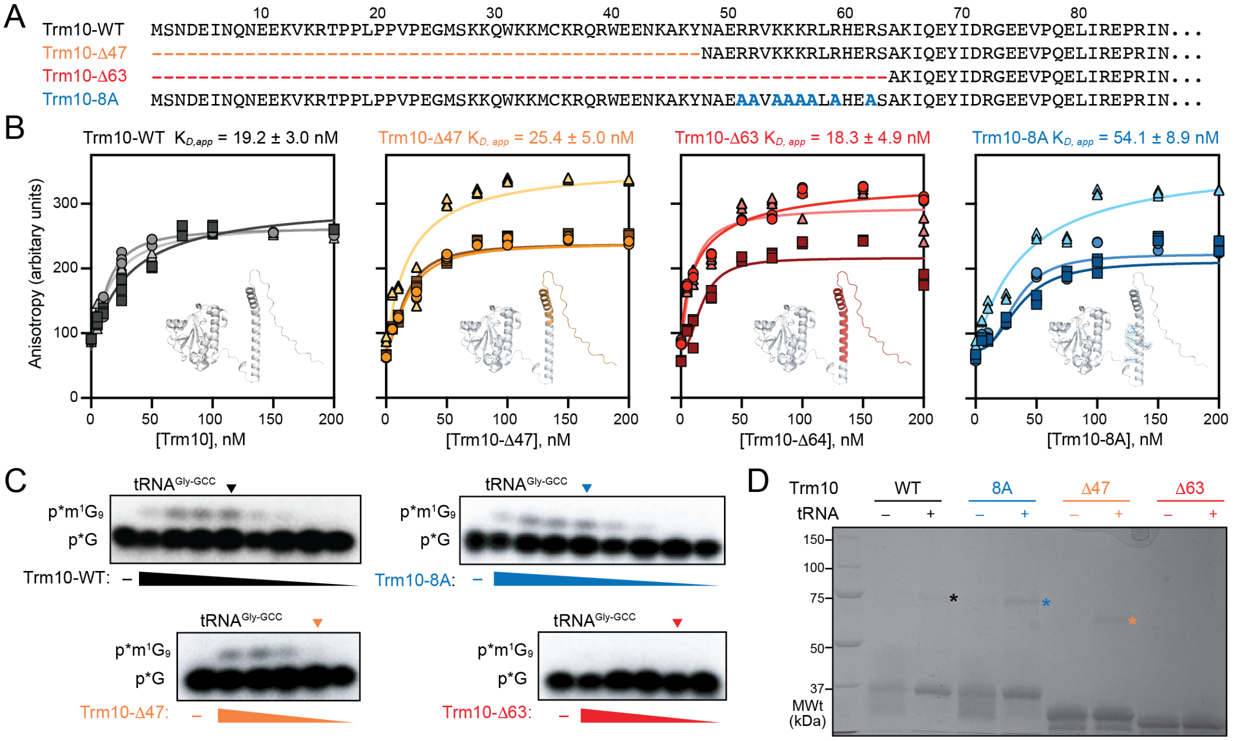


**Fig. S6. The yeast Trm10 NTD does not contribute to tRNA binding and only a limited region plays a role in activity.** ***A,*** Sequence alignment of wild-type Trm10 (Trm10-WT) and NTD variants showing the sites of truncations (Trm10-Δ47 and Trm10-Δ63) and amino acid substitutions in the multisite R/K to A variant (Trm10-8A). ***B,*** Fluorescence anisotropy binding affinity analyses of WT and variant Trm10 proteins to tRNA$^{Gly\text{-}GCC}$ shown for three independent assays that were individually fit to **eq. 1** to yield $K_{D,app}$ and averaged to give the value reported above each plot (see Methods for details). Dotted lines correspond to the WT average shown for reference. Colored regions on the *inset* Trm10 AlphaFold model indicate the locations of deleted or altered residues compared to WT. ***C,*** *In vitro* methyltransferase activity assays using WT and variant Trm10 proteins with uniformly $^{32}$P-G labeled tRNA$^{Gly\text{-}GCC}$ and the indicated enzymes. Wedges shown below each activity panel indicate 5-fold serial dilutions, starting from 7.7 μM Trm10-WT and Trm10-8A, 9.0 μM Trm10-Δ47 and 9.5 μM Trm10-Δ63, with no enzyme lanes indicated "-". Arrowheads above each activity panel mark the dilution that corresponds to ~0.1 μM concentration enzyme in each assay. Assays were performed in duplicate, with a representative assay used for quantification as described in the text shown here. ***D,*** BS3-mediated crosslinking studies of the WT and variant Trm10 proteins were performed as shown in **Fig 6G**, and asterisks indicate tRNA-dependent higher molecular weight bands that correlate with dimer formation for WT and two of the three tested variants. Analyses shown were performed in the absence of SAM; essentially identical results were obtained with the cosubstrate included in the reaction.

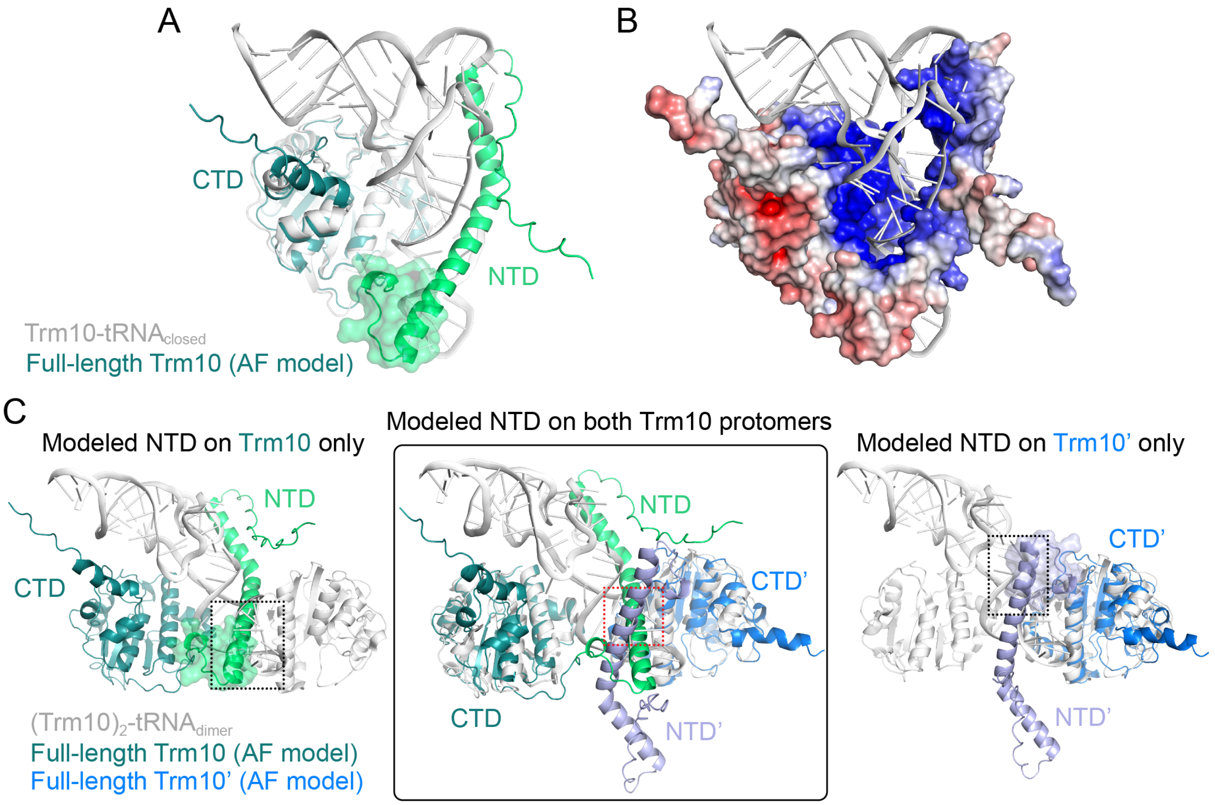


**Fig. S7. AlphaFold (AF) model of full-length *S. cerevisiae* Trm10.** ***A,*** Comparison of the AF model of Trm10 with the Trm10-tRNA$_{closed}$ complex reveals the potential for the Trm10 NTD to wrap around the tRNA. The NTD region proximal to the SPOUT domain that is important for tRNA modification is highlighted with semi-transparent surface, adjacent to the tRNA anticodon stem-loop. ***B,*** Modeling of the NTD-tRNA interaction extends the positive surface of Trm10 engaged with the tRNA. ***C,*** Alignments of the full-length Trm10 AF model on the (Trm10)$_2$-tRNA complex shown for alignment with Trm10 only (*left*), Trm10' only (*right*), and both protomers (*center*). For both modeled full-length proteins, the NTD region proximal to the SPOUT domain (semi-transparent surfaces in *left* and *right* images) are positioned adjacent to each other and the tRNA anti-codon stem-loop (dotted black line boxes) and no major clashes with the experimentally positioned SPOUT domain of the other protomer are observed. When both modeled proteins are aligned on the structure, a clash is observed for their extended N-terminal structure (red dotted line box).

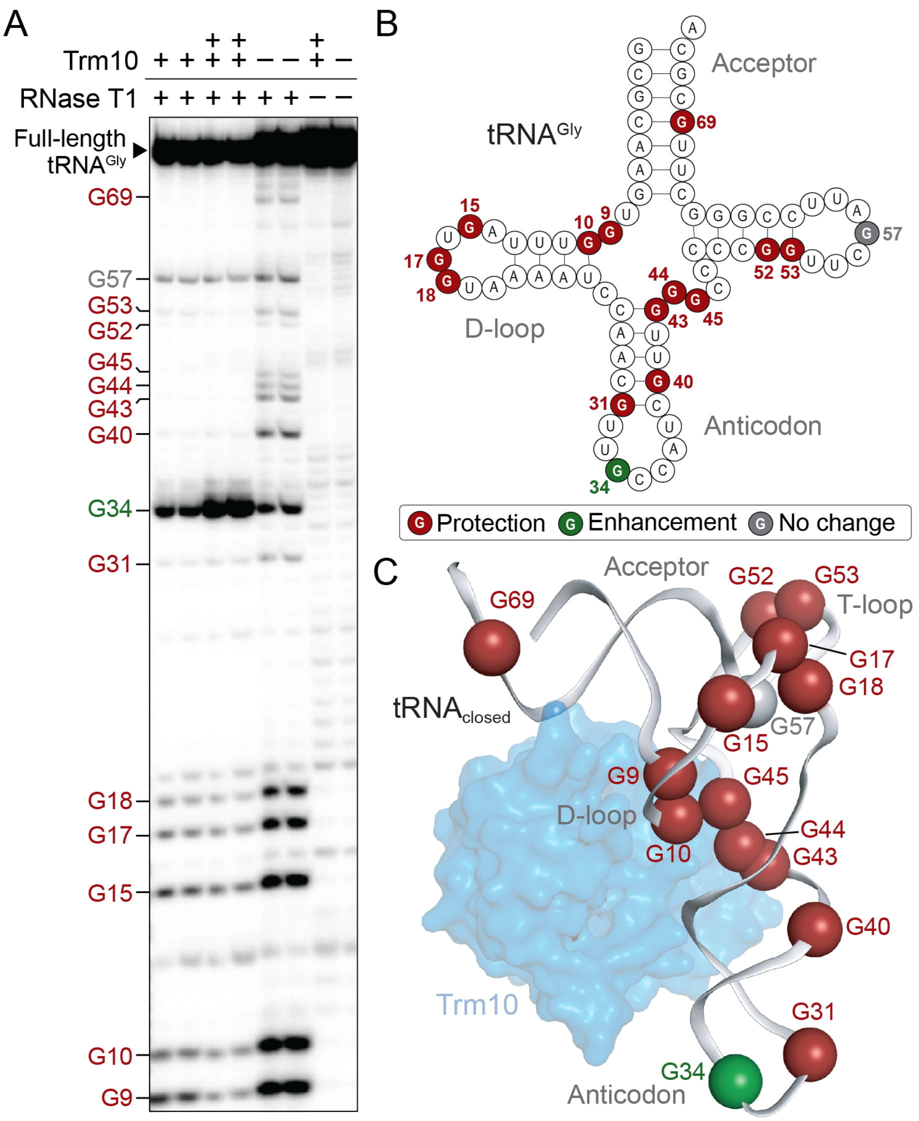


**Fig. S8. RNase T1 nuclease footprinting of the *S. cerevisiae* Trm10-tRNA$^{Gly}$ complex. *A,*** RNase T1 nuclease footprinting analysis of $^{32}$P-5’-end labeled tRNA$^{Gly}$ in the presence or absence of Trm10, shown in duplicate at each protein (1 and 5 μM are indicated + and ++, respectively; no protein is indicated –). The right two lanes show controls without nuclease, both with and without Trm10. Bands were identified based on the known sequence and alkaline hydrolysis ladder (not shown). Footprinting pattern−protection, enhancement, or no change–are mapped on ***B,*** the tRNA$^{Gly}$ secondary structure, and ***C,*** the tRNA$^{Gly}$ from the monomeric Trm10-tRNA$_{closed}$ complex.

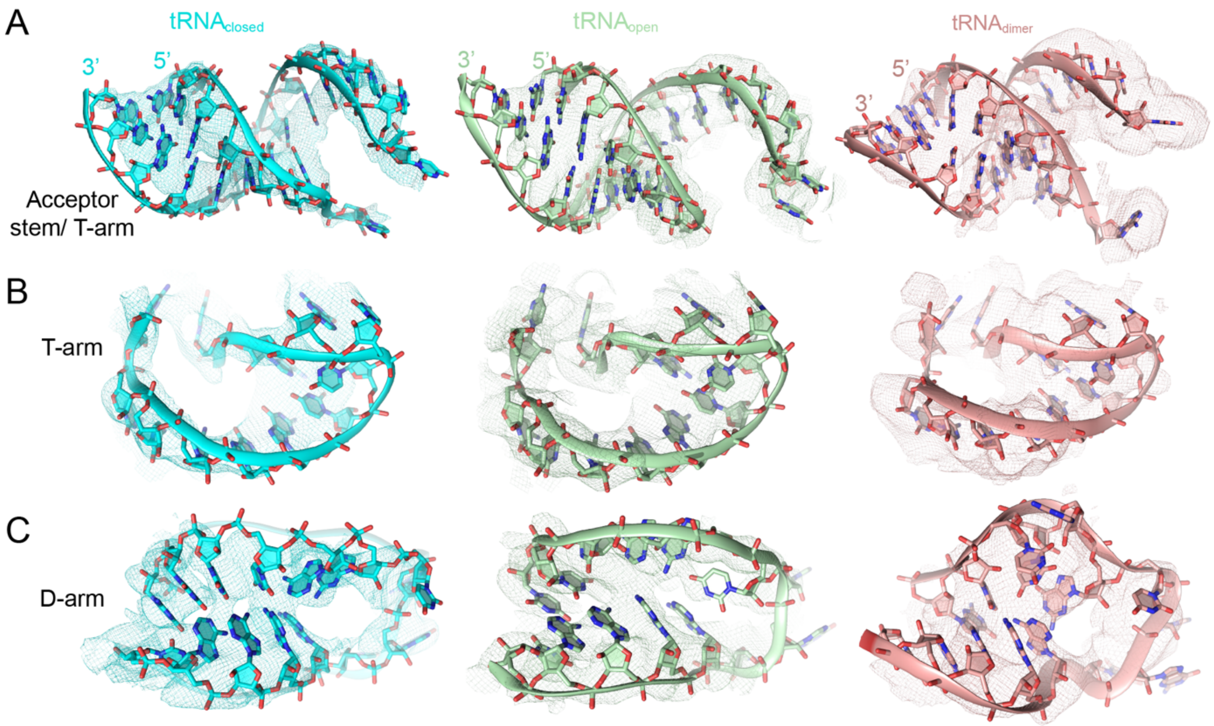


**Fig. S9. Model in cryo-EM map for the tRNA acceptor stem, T-, and D-arm in each Trm10-tRNA complex**. tRNA regions indicated are shown within the final complete map (mesh) with residues and maps are colored as in the main text figures. Model and map (*left to right*) of ***A*,** the acceptor stem/ T-arm from $tRNA_{closed}$ (DEM-sharpened, threshold: 0.458), $tRNA_{open}$ (CS-sharpened; threshold: 0.044), and the dimeric Trm10-tRNA complex ($tRNA_{dimer}$**)** (CS-sharpened; threshold: 0.019). ***B*,** T-arm from $tRNA_{closed}$ (CS-sharpened; threshold: 0.045), $tRNA_{open}$ (CS-sharpened, threshold: 0.033), and $tRNA_{dimer}$ (CS-sharpened; threshold: 0.015). ***C*,** D-arm from $tRNA_{closed}$ (CS-sharpened; threshold: 0.039), $tRNA_{open}$ (CS-sharpened; threshold: 0.057), and $tRNA_{dimer}$ (CS-sharpened; threshold: 0.028). The range of CS-sharpened maps for $tRNA_{closed}$, $tRNA_{open}$, and $tRNA_{dimer}$ are -0.253–0.335, -0.402–0.52, and -0.0897–0.164, respectively. For $tRNA_{closed}$, the range of DEM-sharpened map is -0.0017–2.00.

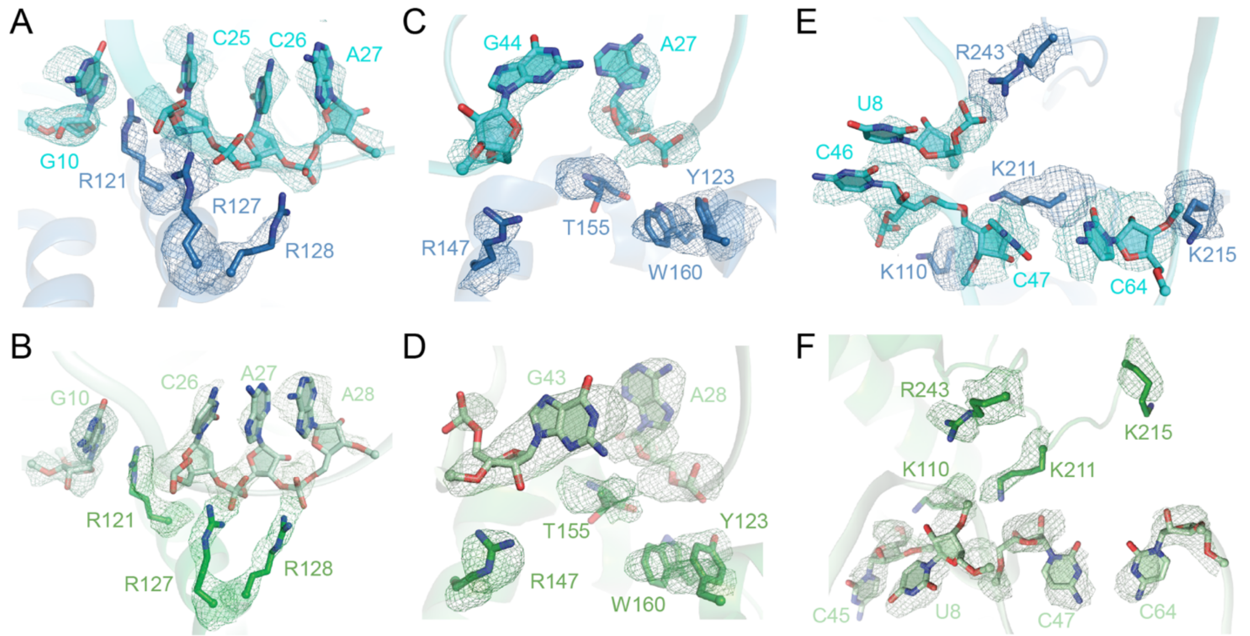


**Fig. S10. Cryo-EM maps of the residues interacting with the tRNA in the monomeric Trm10-tRNA complexes.** Amino acid residues and nucleotides discussed in the main text are shown with the final complete map (mesh) from the two Trm10-tRNA complex structures. The residues and maps are colored as in the main text figures. The range of CS- and DEM-sharpened maps is as noted for *Fig. S7*. For $tRNA_{open}$, the range of DEM-sharpened maps is -0.00338-2.19. Model and map of ***A-B***, R121 ($tRNA_{closed}$, DEM, threshold: 0.34; $tRNA_{open}$, CS, 0.134), R127 ($tRNA_{closed}$, CS, threshold: 0.034; $tRNA_{open}$, DEM, 0.37), R128 ($tRNA_{closed}$, CS, threshold: 0.034; $tRNA_{open}$, DEM, 0.247), G10 ($tRNA_{closed}$, CS, threshold: 0.047; $tRNA_{open}$, CS, 0.088), C25-A27 ($tRNA_{closed}$, CS, threshold: 0.078), C26 ($tRNA_{open}$, CS, threshold: 0.087), A27 ($tRNA_{open}$, CS, threshold: 0.083), A28 ($tRNA_{open}$, unsharpened, threshold: 0.038, range:-0.0463-0.0954). ***C-D,*** R147 ($tRNA_{closed}$, DEM, threshold: 0.281; $tRNA_{open}$, DEM, threshold: 0.21), T155 ($tRNA_{closed}$, DEM, threshold: 0.281; $tRNA_{open}$, DEM, threshold: 0.277), W160 ($tRNA_{closed}$, CS, threshold: 0.099; $tRNA_{open}$, DEM, threshold: 0.543), Y123 ($tRNA_{closed}$, DEM, threshold: 0.524; $tRNA_{open}$, CS, threshold: 0.151), G44 ($tRNA_{closed}$, DEM, threshold: 0.75), G43 ($tRNA_{open}$, DEM, threshold: 0.271), A27 ($tRNA_{closed}$, CS, threshold: 0.082), A28 ($tRNA_{open}$, DEM, threshold: 0.203). ***E-F,*** K110 ($tRNA_{closed}$, CS, threshold: 0.02; $tRNA_{open}$, CS, threshold: 0.06), K211 ($tRNA_{closed}$, DEM, threshold: 0.066; $tRNA_{open}$, CS, threshold: 0.084), K215 ($tRNA_{closed}$, CS, threshold: 0.007; $tRNA_{open}$, DEM, threshold: 0.21), R243 ($tRNA_{closed}$, CS, threshold: 0.019; $tRNA_{open}$, CS, threshold: 0.024), C46 ($tRNA_{closed}$, DEM, threshold: 0.362), C45 ($tRNA_{open}$, CS, threshold: 0.081), U8 ($tRNA_{closed}$, DEM, threshold: 0.244; $tRNA_{open}$, CS, threshold: 0.045), C47 ($tRNA_{closed}$, DEM, threshold: 0.628; $tRNA_{open}$, CS, threshold: 0.077), C64 ($tRNA_{closed}$, DEM, threshold: 0.495; $tRNA_{open}$, DEM, threshold: 0.469).

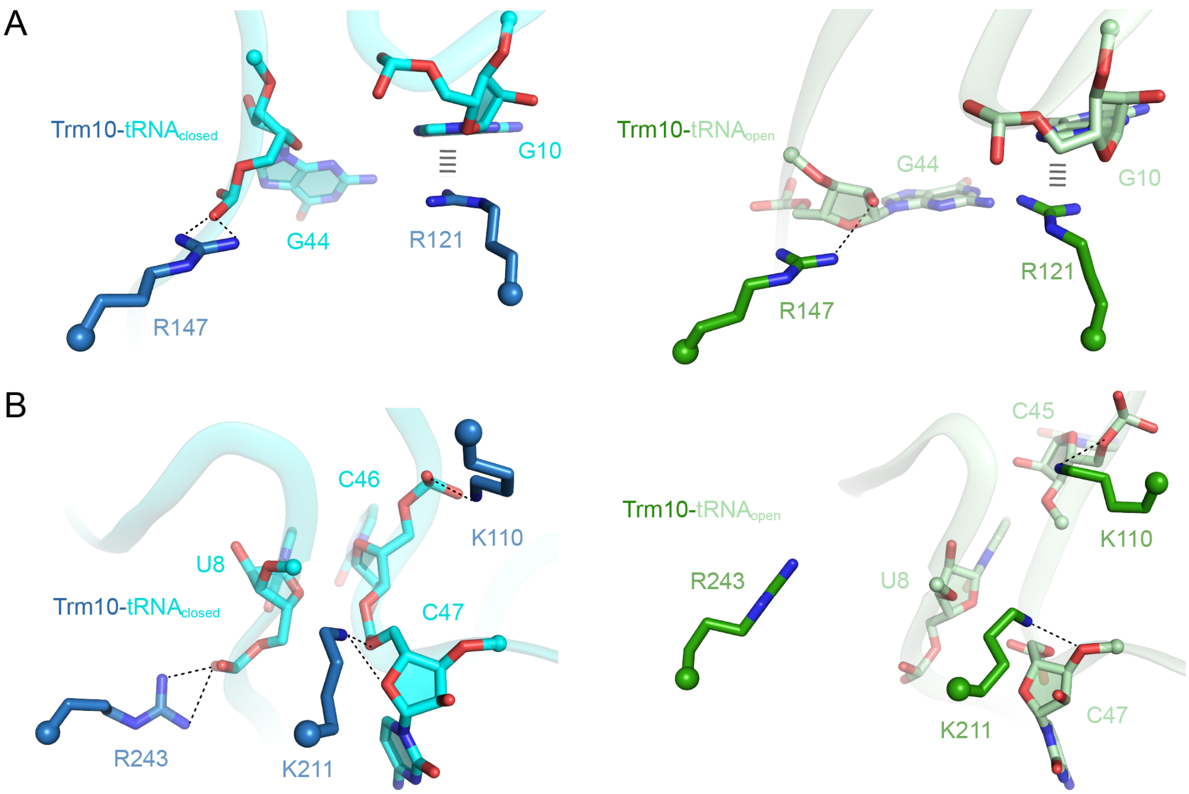


**Fig. S11. Structural features of the Trm10-tRNA$_{closed}$ and Trm10-tRNA$_{open}$ complexes.** ***A,*** At the Trm10-tRNA interface, residues R121 and R147 act as a molecular pincer to secure the two opposing tRNA strands by contacting the phosphate backbone of G44 (dotted black lines) and forming cation-π interactions (gray dashed line) with G10, respectively, in both the tRNA$_{closed}$ and tRNA$_{open}$ conformation. ***B,*** Contacts to both strands at a second site made by K110, K211, and R243, which contact the backbone of C46, C47, and U8, respectively, additionally support RNA-protein interaction in the tRNA$_{closed}$ conformation (*left*). In tRNA$_{open}$, this second set of interactions is only partially present as the tRNA stand rotates away from the enzyme, breaking the interaction with R243 (*right*).

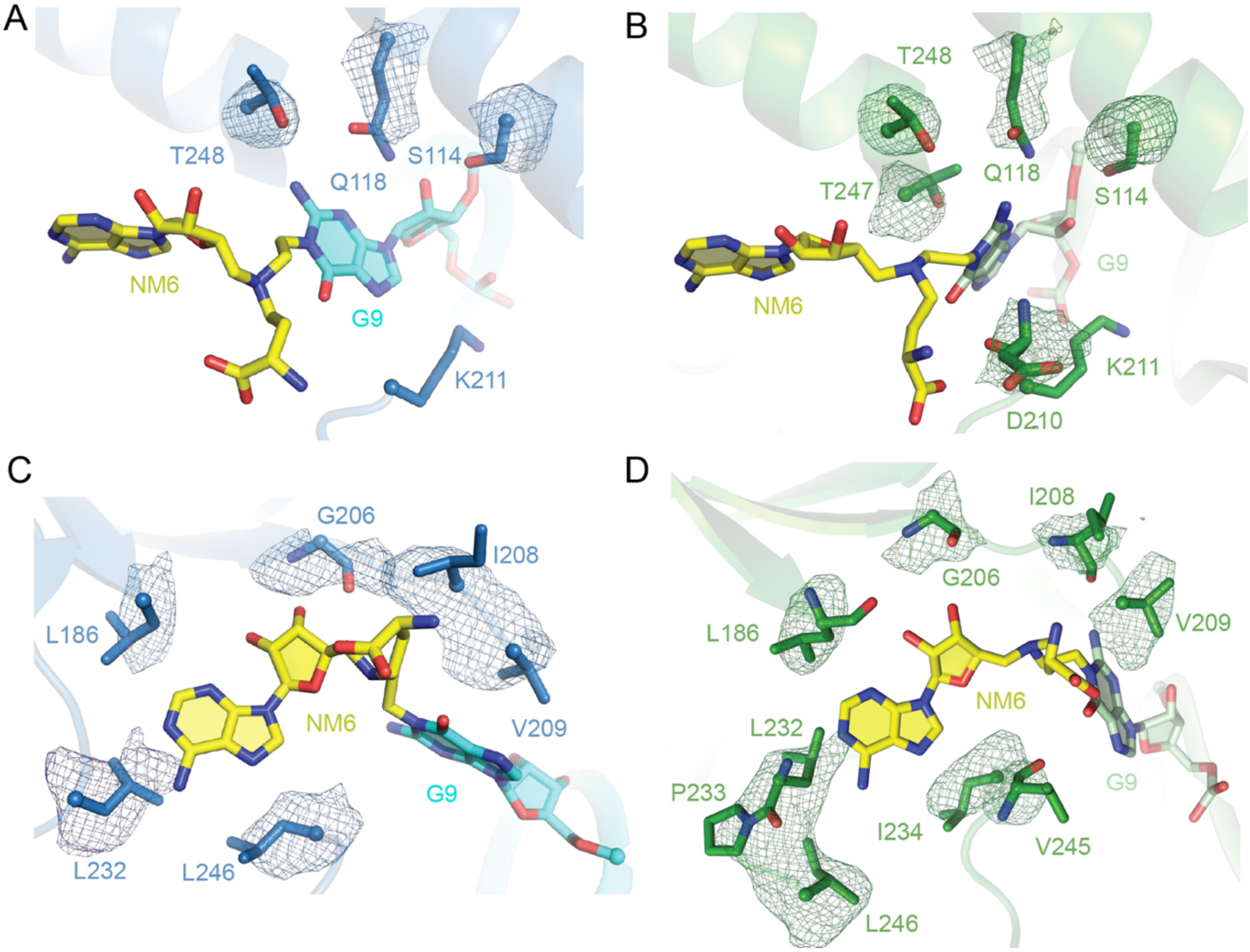


**Fig. S12. Cryo-EM maps corresponding to residues interacting with the covalently linked G9 and NM6 in the monomeric Trm10-tRNA complexes.** Amino acid residues and nucleotides discussed in the main text are shown with the final complete map (mesh). The residues and maps are colored as in the main text figures. The range of CS- and DEM-sharpened maps is as noted in *Fig. S7*. Model and map of ***A-B,*** T248 ($tRNA_{closed}$, DEM, threshold: 0.753; $tRNA_{open}$, DEM, threshold: 0.58), Q118 ($tRNA_{closed}$, CS, threshold: 0.12; $tRNA_{open}$, DEM, threshold: 0.165), S114 ($tRNA_{closed}$, DEM, threshold: 0.459; $tRNA_{open}$, DEM, threshold: 0.444), T247 ($tRNA_{open}$, DEM, threshold: 0.314), D210 ($tRNA_{open}$, CS, threshold: 0.065). ***C-D,*** L186 ($tRNA_{closed}$, DEM, threshold: 0.5; $tRNA_{open}$, DEM, threshold: 0.339), G206 ($tRNA_{closed}$, DEM, threshold: 0.443; $tRNA_{open}$, DEM, threshold: 0.481), I208 ($tRNA_{closed}$, CS, threshold: 0.066; $tRNA_{open}$, CS, threshold: 0.08), V209 ($tRNA_{closed}$, CS, threshold: 0.066; $tRNA_{open}$, DEM, threshold: 0.31), L232 ($tRNA_{closed}$, DEM, threshold: 0.111; $tRNA_{open}$, DEM, threshold: 0.265), L246 ($tRNA_{closed}$, CS, threshold: 0.047; $tRNA_{open}$, DEM, threshold: 0.253), V245 ($tRNA_{open}$, DEM, threshold: 0.117), L232, P233, and I234 ($tRNA_{open}$, DEM, threshold: 0.265).

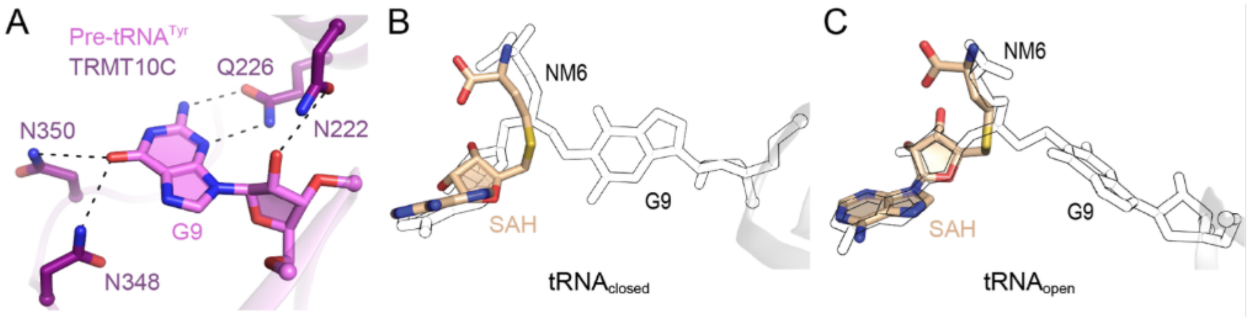


**Fig. S13. Recognition of G9 in human TRMT10C and mimicry of SAH by NM6 in Trm10-tRNA complexes.** ***A,*** In the hTRMT10C-pre-tRNA$^{Tyr}$ complex (PDB 7ONU), N222 engages the ribose sugar, Q226 recognizes both the nucleobase N3 and primary amine group (exocyclic N2), while N348 and N350 coordinate with the hydroxyl group (exocyclic O6) of the G9 target nucleotide. ***B-C,*** NM6 bound to G9 (outline) is in a bent conformation matching the pose observed for SAH (yellow) in the *S. cerevisiae* Trm10 structure (PDB 4JWJ).

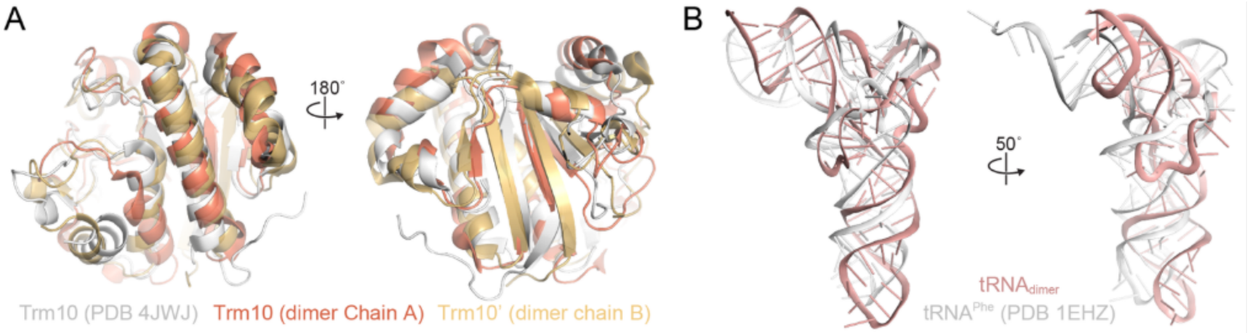


**Fig. S14. Comparison of the (Trm10)$_2$-tRNA complex with free Trm10 and tRNA.** ***A,*** Structural alignment of the two Trm10 protomers from the dimeric complex with free Trm10 (PDB 4JWJ) shows that the overall structure remains largely unchanged. ***B,*** Superimposition of the tRNA from the (Trm10)$_2$-tRNA complex with unbound tRNA$^{Phe}$ (PDB 1EHZ) reveals pronounced differences in multiple regions.

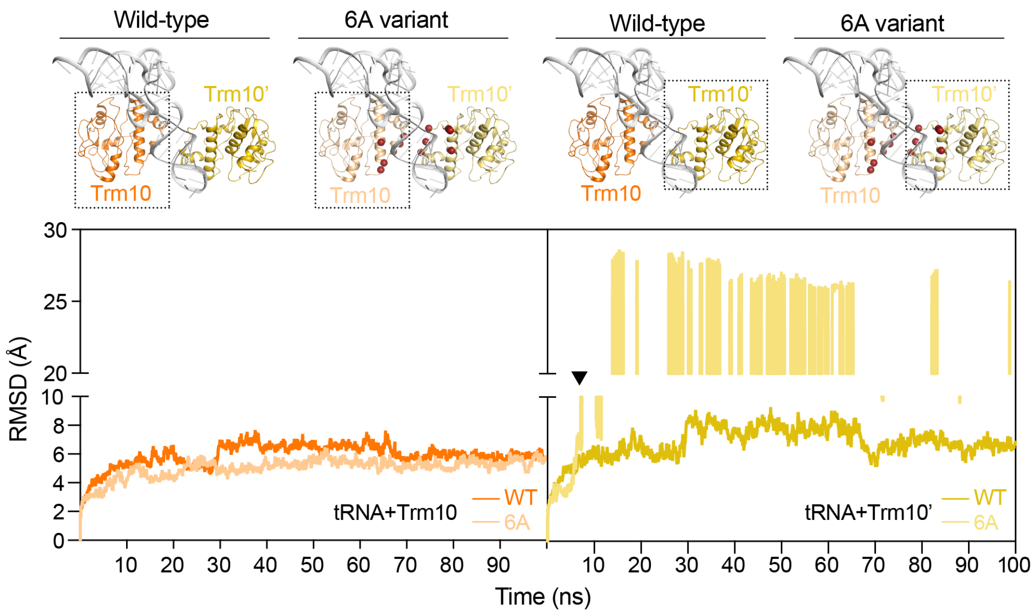


**Fig. S15. MD simulation of the (Trm10)$_2$-tRNA complex: both protomers as the 6A variant.** Plots of system RMSD for the indicated Trm10 protomer (Trm10 *left* and Trm10' *right*) and tRNA chains. Cartoons of the simulated systems (*top*) highlight the protomer-tRNA complex analyzed below (boxed) and the locations of the 6A substitutions in both protomer of the 6A variant (red spheres: R121A/ R127A/ R128A/ N154A/ N156A/ N159A). The mutated Trm10' protomers dissociates early in the simulation (*right* plot, indicated by the arrowhead) and as a result no further analyses were performed on this complex.

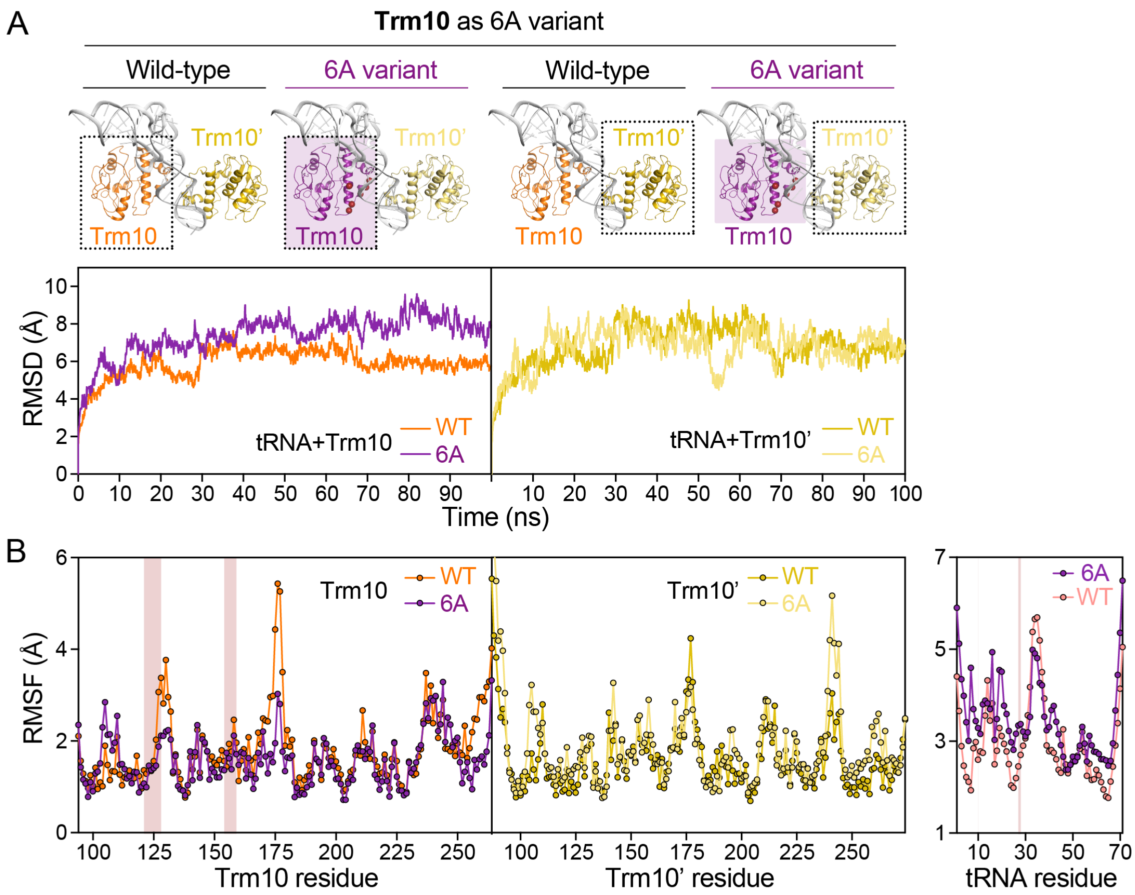


**Fig. S16. MD simulation of the (Trm10)$_2$-tRNA complex: Trm10 protomer as the 6A variant.** ***A,*** Plots of system RMSD for the indicated Trm10 protomer (Trm10 *left* and Trm10' *right*) and tRNA chains. Cartoons of the simulated systems (*top*) highlight the protomer-tRNA complex analyzed below (boxed) and the locations of the 6A substitutions in Trm10 protomer only (red spheres: R121A/ R127A/ R128A/ N154A/ N156A/ N159A). ***B,*** Plots of residue RMSF for the Trm10 (*left*) and Trm10' (*center*) protomers, and the tRNA (*right*). The data shown in these plots were used to calculate ΔRMSF values (6A variant-containing minus wild-type) shown in **Fig. 7A**.

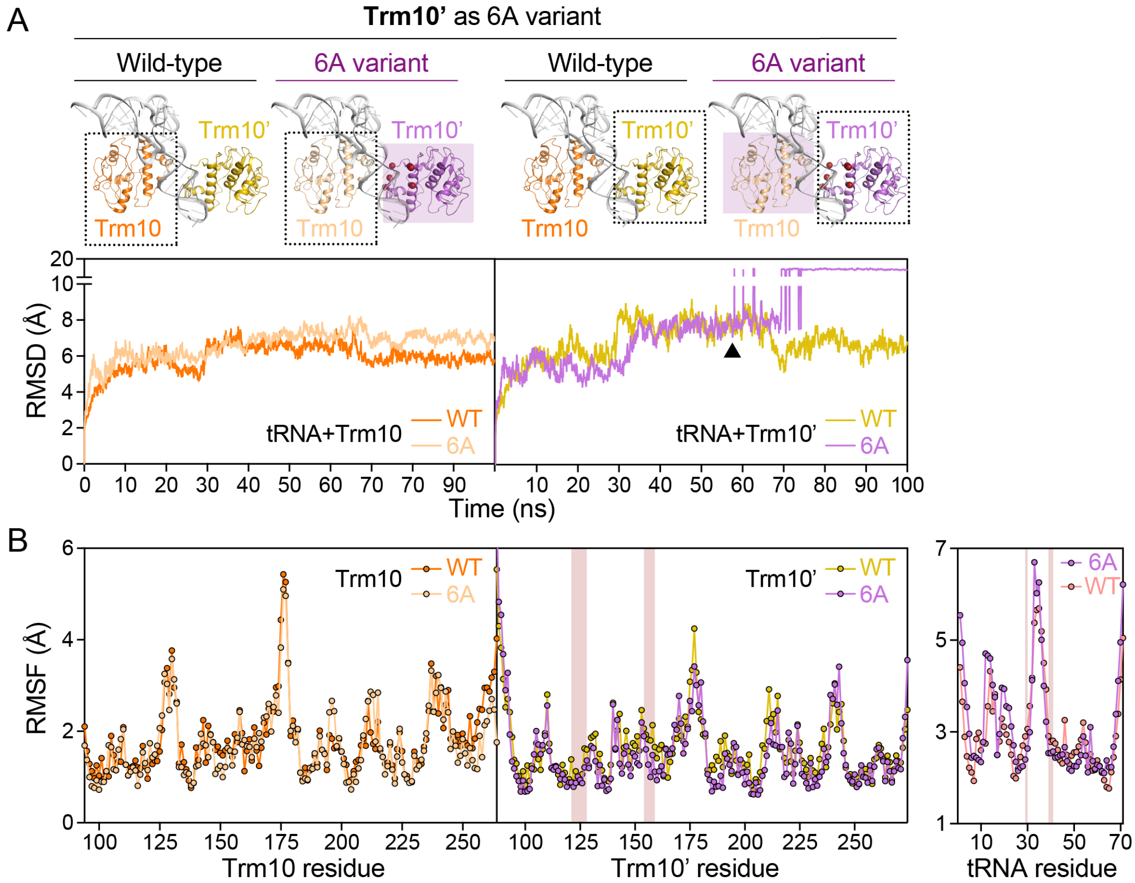


**Fig. S17. MD simulation of the (Trm10)$_2$-tRNA complex: Trm10' protomer as the 6A variant.** ***A,*** Plots of system RMSD for the indicated Trm10 protomer (Trm10 *left* and Trm10' *right*) and tRNA chains. Cartoons of the simulated systems (*top*) highlight the protomer-tRNA complex analyzed below (boxed) and the locations of the 6A substitutions in Trm10' protomer only (red spheres: R121A/ R127A/ R128A/ N154A/ N156A/ N159A). ***B,*** Plots of residue RMSF for the Trm10 (*left*) and Trm10' (*center*) protomers, and the tRNA (*right*). The data shown in these plots were used to calculate ΔRMSF values (6A variant-containing minus wild-type) shown in **Fig. 7B**.

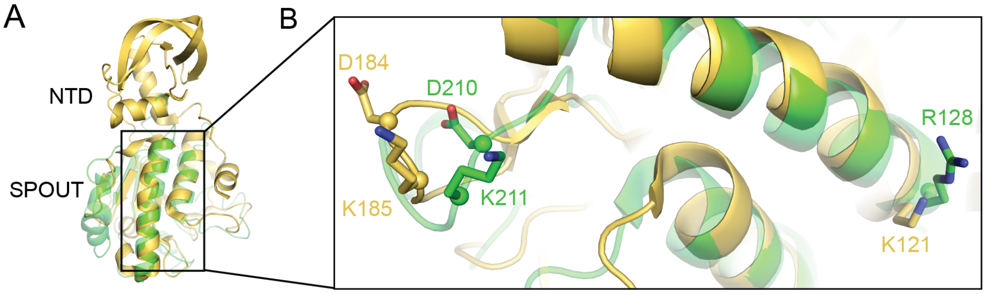


**Fig. S18. Comparison of *S. cerevisiae* and *S. acidocaldarius* Trm10 proteins**. ***A***, Alignment of the *S. cerevisiae* Trm10 SPOUT domain from the tRNA$_{open}$ complex (green) and *S. acidocaldarius* Trm10 (PDB 5A7T). ***B***, Zoomed in view highlighting the alignment of key residues in the two structures, as discussed in the main text.

### SUPPLEMENTARY TABLES

**Table S1. Amino acid sequence conservation among fungal and human Trm10 enzymes**

| *S. cerevisiae* Trm10 | All fungi (% occurrence) | Human TRMT | | |
|---|---|---|---|---|
| | | 10A | 10B | 10C |
| K110 | K (62.5), N (11.1) | K | K | K |
| S114 | S (97.7) | K | R | N |
| Q118 | Q (98.5) | Q | Q | Q |
| R121 | R (76.5), Y (12.9) | R | R | E |
| Y123 | Y (95.8) | Y | Y | E |
| R127 | R (61.5), K (25.3) | R | K | R |
| R128 | R (14.8); NC[a] | R | K | R |
| R147 | R (79.2) | K | Y | H |
| T155 | T (4.2); NC[a] | K | D | Q |
| W160 | W (95.1) | W | Y | W |
| L186 | L (98.9) | L | L | L |
| G206 | G (100) | G | G | G |
| I208 | I (64.0), L (33.1) | L | L | F |
| V209 | V (90.8) | V | V | V |
| D210 | D (100) | D | D | D |
| K211 | K (64.7), R (25.0) | H | E | K |
| K215 | K (95.2) | K | K | P |
| L232 | L (100) | L | L | L |
| P233 | P (99.6) | P | P | P |
| I234 | I (97.8) | L | I | L |
| R243 | R (98.3) | R | K | K |
| V245 | V (93.3) | V | I | - |
| L246 | L (97.0) | L | L | L |
| T247 | T (65.1), A (30.0) | A | A | T |
| T248 | T (54.3), V (37.9) | V | I | L |

[a]Not conserved
